# Rethinking scRNA-seq Trajectories in Phylogenetic Paradigms: Overcoming Challenges of Missing Ancestral Information

**DOI:** 10.1101/2025.07.22.664676

**Authors:** Julia Naas, Arndt von Haeseler, Christiane Elgert

**Author notes:** Correspondence: Julia Naas.

## Abstract

In recent decades, many bioinformatics tools have been developed to reconstruct trajectories of biological processes, e.g., cell differentiation, using single-cell RNA-sequencing (scRNA-seq) data. Most tools tacitly assume that a cell’s ancestral transcriptomic profile can be approximated by means of its neighboring cells in an embedded gene expression space. However, many scRNA-seq datasets lack ancestral information due to missing early or transient states at the time of sequencing. We introduce CellREST, a bioinformatics tool that reformulates trajectory reconstruction as a phylogenetic inference problem. It infers trees linking cells that are assumed to share a common ancestral expression state. Using maximum likelihood tree inference, CellREST uncovers multiple different aspects of the transcriptomic landscape underlying a single scRNA-seq dataset, which can be visualized and combined into a single-cell network. We showcase CellREST’s performance on simulated and experimental scRNA-seq data and recover circular processes as well as cell type converging differentiation scenarios. By introducing and adapting phylogenetic concepts, CellREST provides a framework for interpreting transcriptomic relationships between cells within scRNA-seq data.

## Introduction

Single-cell RNA sequencing (scRNA-seq) provides indepth insights into transcriptomic profiles of individual cells, enabling detailed analysis of heterogeneity within and between different cell types. However, one scRNAseq dataset captures gene expression only as a snapshot, limiting the direct observation of ongoing or past dynamical processes, e.g., cell type differentiation.

To infer the transcriptomic history of a cell many bioinformatics methods have been developed, for example Monocle1-3 (1–3), Slingshot (4), Palantir (5), STREAM (6), CellRank1 and 2 (7, 8) or scFates (9). While extensive efforts have been made to evaluate the performance and applicability of such tools (10), most rely on approximating a cell’s ancestral gene expression profile using those of its neighbors within method-specific embeddings. This approach implicitly assumes that transcriptomically similar cells follow the same differentiation trajectory, which is a reasonable premise when the process is continuously and evenly sampled. However, in practice, early or transient states are often underrepresented due to rapid progression or limited sampling. In such cases, embedding-based distance measures may fail to relate cells with shared transcriptomic histories if they appear distant in the chosen embedding space.

This key limitation of scRNA-seq data, namely the lack of explicit ancestral information, has long been addressed in phylogenomics. Phylogenomics aims to infer evolutionary relationships between biological entities, typically species, by analyzing observed heritable traits, such as morphological characteristics or molecular sequences. Relationships are typically visualized as phylogenetic trees, grouping species to indicate a shared common ancestor.

There are two principal approaches to infer a phylogenetic tree. *Distance-based methods*, as for example Neighbor Joining (NJ) (11) or BIONJ (12), construct trees by grouping species based on their pairwise distances. In contrast, *character-based methods*, such as Maximum Likelihood (ML) (13), model the evolution of individual character states, for example, nucleotide bases A, C, G, and T. Maximum Likelihood infers the tree that best explains the observed data, where branch lengths represent the extent of genetic change over time.

Tree reconstruction methods based on distances of expression profiles of cells or (pseudo-)bulks have already been used to analyze transcriptomic data (14–16). However, like the above mentioned trajectory reconstruction methods, they share the limitation to depend on predefined distance or similarity measures. More recently, also ML-based (17) and Brownian motion–based tree reconstruction methods (18) have been applied to scRNA-seq data. These available methods either compare transcriptomic trees to those derived from single-nucleotide polymorphisms (SNPs) (17), or mix information on the evolution of species with transcriptomic changes (18). While species evolution, cell division into lineages, and transcriptomic changes are undoubtedly interdependent (19, 20), we argue that these processes cannot be fully represented within a single tree. Furthermore, methods that rely solely on a single tree structure are insufficient to capture nontree-like features of the transcriptomic landscape, i.e., the manifold formed by gene expression profiles of cells undergoing a biological process.

To close this gap, we introduce **CellREST** (**Cell**ular **R**elationships in **E**xpression-state-based **S**ingle-cell **T**rees), a phylogenetic paradigm for the analysis of single-cell transcriptomic data. CellREST uses a character-based phylogenetic framework for inferring gene expression state transitions between cells in scRNAseq data. It reconstructs a collection of single-cell-labeled Maximum Likelihood trees that capture alternative and potentially coexisting trajectories. Inferred trees are combined into a single-cell network that displays complex, non-tree-like transcriptomic landscapes in a noise-robust manner. Implemented as an R package, CellREST integrates with the Seurat workflow (21), allowing further data-driven exploration.

## Results

### A Phylogenetic Paradigm for Exploring Transcriptional Landscapes

To address the problem of analyzing trajectories on scRNA-seq data in phylogenetic terms, the pre-processed, log-normalized count matrix of a scRNAseq dataset (Methods M.1) is subset to highly variable genes and transformed into a pseudo-alignment. For each gene, expression levels are discretized into four categorical expression states: Zero expression values are assigned to a ‘no expression’ state and remaining positive expression values are uniformly divided into three bins representing ‘low expression’, ‘medium expression’, and ‘high expression’ states (Methods M.2.1). Fig. S1A shows that there is no substantial difference between UMAP (22, 23) embeddings for discretized data (in the exemplary downstream analysis encoded as ordered integers 0, 1, 2, and 3) and non-discretized data. This suggests that discretization does not reduce biologically meaningful variation in the data. To enable character-based phylogenetic inference, the four expression states are encoded as pseudo-nucleotides T (no expression), G (low expression), C (medium expression), and A (high expression) (Fig. 1A 1-2). This results in a pseudo-alignment in which each column (‘pseudo-site’) corresponds to a gene and each row represents a cell (‘pseudo-taxon’). Encoding expression states as pseudo-nucleotides does not preserve their low to high ordering, but enables the estimation of transition dynamics through phylogenetic inference.

**Fig. 1.**
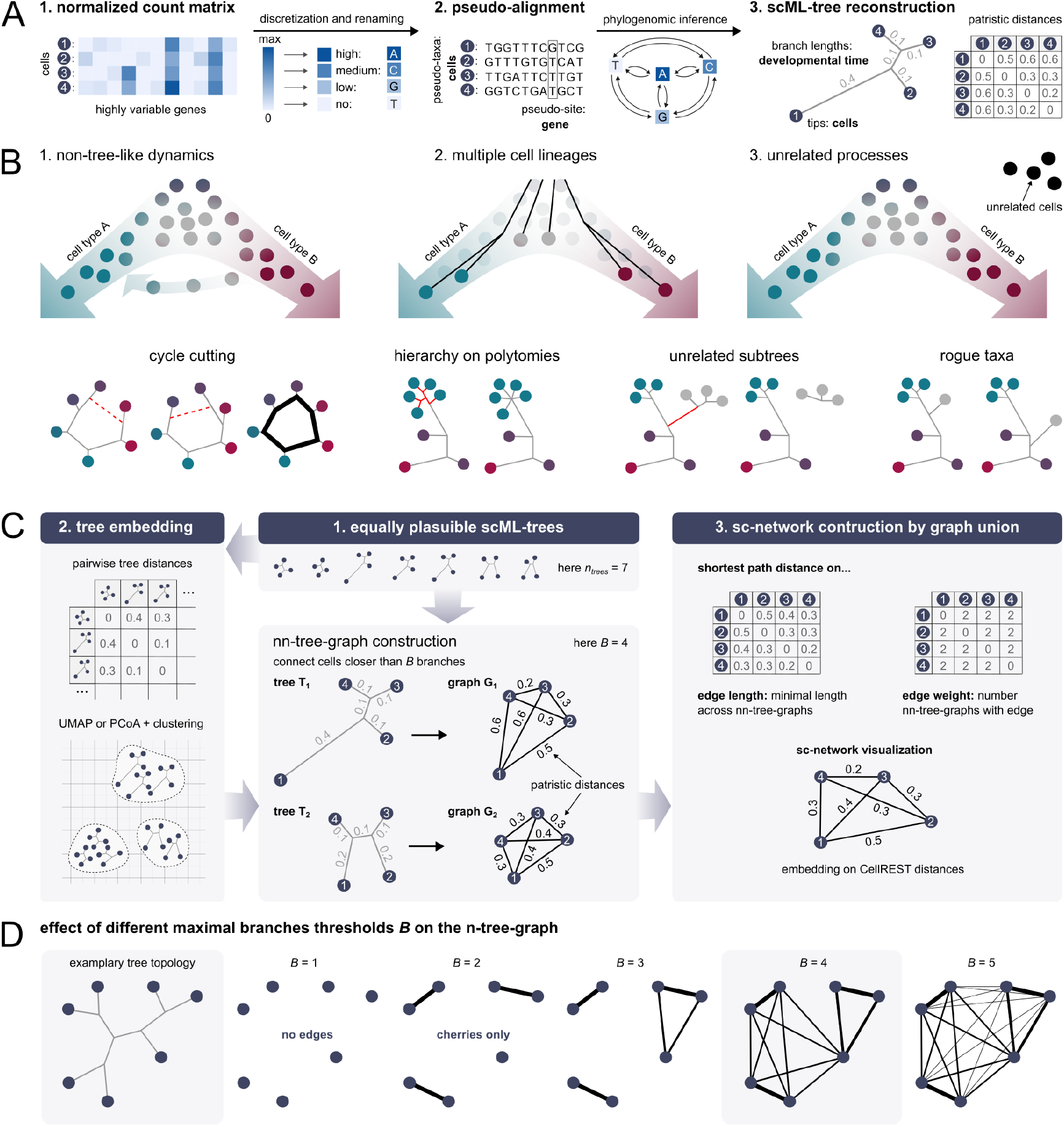
A Phylogenetic view on scRNA-seq data. A CellREST 1. subsets the scRNA-seq log-normalized cell-times-gene matrix to highly variable genes, 2. transforms the resulting data into a pseudo-alignment and 3. infers single-cell-labeled Maximum Likelihood trees (scML-trees). **B** A single tree topology is insufficient to display 1. non-tree-like dynamics, 2. multiple cell lineages, or 3. unrelated processes, leading to phylogenetic challenges such as polytomies and rogue taxa / subtrees. **C** CellREST reconstructs multiple scML-trees and 1. evaluates their variability via pairwise tree-distances and corresponding tree-embeddings, and 2. transforms all or a subset of scML-trees into nn-tree-graphs and combines them into a unified single-cell network (sc-network). The sc-network defines the CellREST distance as the shortest path-distance between cells given the edge lengths of the network. **D** Visualization of different *B* thresholds used in constructing nn-tree-graphs.

A phylogenetic tree is inferred from the pseudo-alignment (Methods M.2.2) using the maximum likelihood approach implemented in IQ-TREE2 (24). The resulting single-cell-labeled Maximum Likelihood tree (scML-tree) represents likely ancestral transcriptomic relationships among individual cells. In this scML-tree, branch lengths reflect the number of inferred expression state changes and thus serves as a proxy for development time (Fig. 1A 3). For any pair of cells, the sum of the branch lengths along the connecting path defines their patristic distance.

Across all datasets analyzed in this study, we observed that inferred average transition rates were highest between neighboring expression states, i.e., transitions from no to low, low to medium, and medium to high expression (Fig. S1B). This pattern suggests that tree inference primarily captures gradual changes in gene expression across cells, while still allowing occasional transitions from, e.g., no to medium expression.

Unlike typical studies of species evolution, single-cell transcriptomic data can introduce additional complexities that can make reliance on a single phylogenetic tree potentially inadequate. These include:

#### 1. Non-tree-like dynamics

Dynamical processes, as e.g. the cell cycle (25) or convergent cell type differentiation, can involve non-tree-like topologies (Fig. 1B 1), which cannot be captured by a single tree. As a result, any tree must introduce an artificial ‘cut’ to represent a cycle. The position of this cut depends on both the data support and the heuristics applied during tree reconstruction, leading different trees to resolve the cycle at different locations. This leads to inconsistent placement of cells – in phylogenetic terms known as rogue taxa – whose placements vary across trees due to limited phylogenetic support.

#### 2. Multiple cell lineages

Multiple cell lineages may follow the same dynamical process and are not distinguishable based on transcriptomic information alone (Fig. 1B 2). Ideally, such cells would be represented as polytomies – unresolved branches reflecting shared, indistinct transcriptomic histories. However, since IQ-TREE returns fully bifurcating trees by default, these relationships are resolved arbitrarily, which can misrepresent the underlying transcriptomic structure.

#### 3. Unrelated processes

A scRNA-seq dataset may contain cells originating from distinct biological processes (Fig. 1B 3). Incorporating these unrelated cells into a single phylogenetic tree can result in rogue taxa or subtrees, often connected by disproportionately long branches.

#### 4. Data sparsity and technical noise

The inherent sparsity of scRNA-seq data that is caused by dropouts and low sequencing depth (26) can affect the inferred tree. In particular, technical zeros are misinterpreted as true biological zeros, introducing artificial expression state transitions. This results in longer branch lengths, especially for branches that are connected to the tips of a tree (Fig. S1C). To account for this effect, we apply Adaptively Thresholded Low-Rank Approximation (ALRA) imputation v(27) to log-normalized count matrices prior to binning (Fig. S1D, Methods M.1), which aims to assign positive expression values to technical zeros. This leads to a reduction in branch lengths overall (Fig. S1E).

To overcome the limitations caused by relying on a single phylogenetic tree, we infer multiple scML-trees per dataset (*n*_*trees*_ = 100 for all analyses). In contrast to deterministic methods such as NJ, repeated ML tree inference can recover alternative topologies: This diversity arises from the stochasticity of the tree search, which may converge on different, equally plausible trees depending on initial conditions and local optima in the likelihood space. This is particularly relevant when the data reflect non-tree-like trajectories, where no single optimal topology can fully capture the biological complexity. Each tree thus captures different aspects of the underlying transcriptomic landscape, offering a more nuanced representation of uncertainty in single-cell transcriptomic data.

To explore both the variability across different scML-trees and the complementary information they provide, we consider two approaches:

In the first approach (Fig. 1C 2), we quantify similarities and differences between pairs of scML-trees using a path-based correlation measure (28), which we will refer to as *patristic-correlation* (Methods M.2.3). This measure captures how similarly two trees encode distances between cells. The resulting tree-distance-matrix is visualized using dimensionality reduction techniques such as Principal Coordinates Analysis (PCoA) and UMAP (29–32). In these tree-embeddings, scML-trees with similar topologies and branch lengths cluster together, revealing how many scML-trees support similar parts of the underlying transcriptomic landscape (Fig. 1C 2).

In the second approach (Fig. 1C 3), we use a nearest neighbor (nn) strategy to construct a noise-robust representation of a collection of scML-trees. To do so, each scML-tree is transformed into an nn-tree-graph by connecting cells separated by at most *B* branches (Methods M.2.4). For example, if *B* = 2, only tips that share a direct ancestor are connected (cherries). To capture slightly broader local structures as links between adjacent cherries we use *B* = 4 in all analyses presented here (Fig. 1D). For a collection of scML-trees respective nn-tree-graphs are combined into a single-cell network (sc-network) by including any edge that appears in at least one of the nntree-graphs (Methods M.2.4). Each edge in the sc-network is assigned the following attributes: The **edge weight** reflects how often the edge occurred across the nn-tree-graphs and can be interpreted as a stability score. The **edge length** is defined as the minimum patristic distance between cells that are connected via an edge across all nn-tree-graphs. The edge length hence represents the extent of tree-informed changes in expression profiles. Alternatively, users may choose other distances, e.g., the count of branches between cells, or different summary statistics than the minimum for the attribution of edge lengths.

We define **CellREST distance** as the shortest path distance between cells in the sc-network, measured using edge lengths. This distance metric offers two key advantages over a conventional Euclidean distance between embedded expression profiles: (1) it relies on model-inferred branch lengths rather than geometric proximity, and (2) as shortest path distance, it more accurately approximates curved trajectories. Both CellREST distances and alternative shortest path distances can serve as inputs for downstream analyses including cell embeddings, clusterings, and other exploratory tasks.

### Recovering Complex Trajectories from Simulated scRNA-seq Data

To assess the ability of CellREST to reconstruct non-tree-like trajectories, we simulated scRNA-seq data following a circular topology, such as it occurs, e.g., for cells in cell cycle with respect to cycling genes (25). To do so, we used dyngen (33), which simulates gene expression dynamics over continuous time scale using Gillespie’s stochastic simulation algorithm (34). For a clear and interpretable comparison to the ground truth, we restricted the simulation to a single circular loop (450 cells) and rescaled the simulation time to the interval [0, 1] (Fig. 2A, Methods M.3.1).

**Fig. 2.**
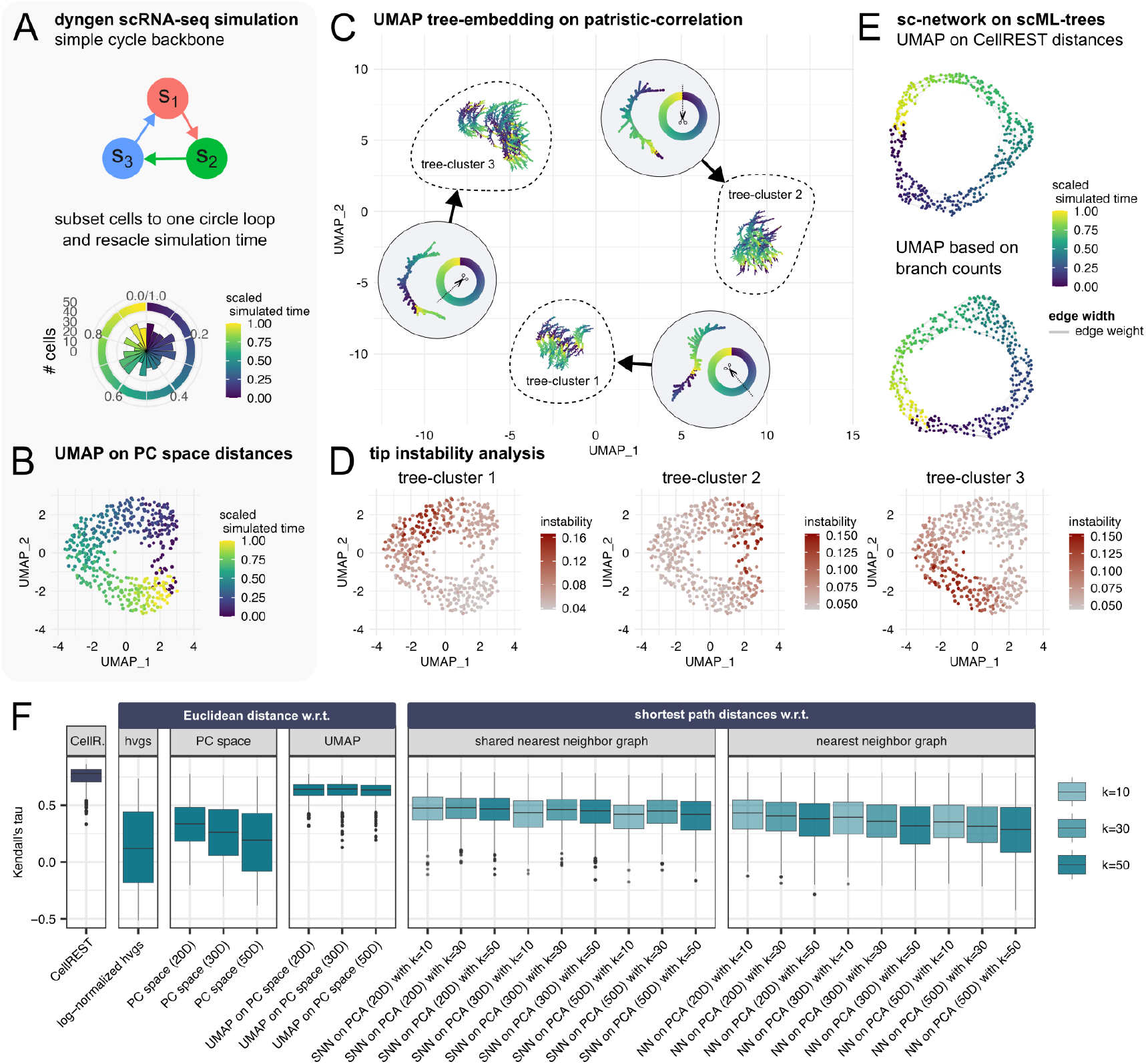
CellREST recovers non-tree-like dynamics in circular trajectories. Abbreviations: dist = distances. **A** Dyngen (33) simulation of a circular differentiation process, restricted to a single loop with simulation time rescaled to the interval [0, 1]. Variation in data support across the trajectory is visible as changes in cell density. **B** Standard UMAP embedding computed on PC space distances. **C** UMAP tree-embedding of 100 scML-trees based on patristic-correlation, with representative tree and cutting location visualized. **D** Tip instability analysis for tree-clusters from panel C. **E** UMAP visualizations of sc-network based on CellREST distances (top) and shortest path distances using minimal branch counts between cells across nn-tree-graphs (bottom); edge widths reflect edge weights. **F** Comparison of Kendall’s tau correlations between simulation time and distances derived from various embeddings, with circular time correction.

Following standard pre-processing and dimensionality reduction using Seurat (Methods M.1), the underlying circular topology was already visible in the UMAP embedding (Fig. 2B). However, we also observed uneven data support across simulation time, with notably lower density in certain regions.

Applying CellREST, we inferred 100 scML-trees, each representing a plausible scenario with similar likelihoods (Fig. S2A, Methods M.2.5). The tree embedding based on patristic-correlation revealed three distinct tree-clusters, each cutting the circular trajectory in a different but consistent region (Fig. 2C). This grouping of scML-trees could not be resolved using splits-based Robinson-Foulds (RF) distances alone (Fig. S2B), highlighting the importance of incorporating branch length information. A tip instability analysis (35), computed within each tree-cluster, enabled localization of the cutting locations through the identification of rogue taxa (see also Fig. 1B 3, Methods M.2.3). These cutting locations coincided with areas of reduced data support (Fig. 2D). When combining all 100 scML-trees into a single sc-network, the full circular trajectory was successfully reconstructed, as visible in a UMAP embedding based on CellREST distances (Fig. 2E). Notably, logarithmized edge lengths in the sc-network closely followed a normal distribution (Fig. S2C, Methods M.2.4).

To quantify how well CellREST distances reflect the simulated ground truth, we conducted a rank-based correlation analysis. Since gene expression changes nonlinearly over simulation time (Fig. S2D), we used Kendall’s tau to compare the relative ordering of cells. For each cell, we ranked all other cells by both simulation time and CellREST distance, accounting for the circular nature of the trajectory, i.e., treating time differences between 0 and 1 as zero (Methods M.3.2). CellREST distances consistently yielded higher Kendall’s tau correlations with simulation time than corresponding analyses with Euclidean distances computed from log-normalized expression profiles or cells in PC space, or UMAP embeddings. Additionally, CellREST outperformed shortest path distances derived from both nearest neighbor (NN) and shared nearest neighbor (SNN) graphs (Fig. S2E, Methods M.1).

### Recovering Missing Transient States from Simulated scRNA-seq Data

Real-world datasets often lack coverage of the entire trajectory. Hence, we further tested CellREST’s robustness by removing cells from two simulation time intervals, *I*_1_ = (0.45, 0.55) and *I*_2_ = (0.15, 0.95), to mimic missing transient states (Fig. 3A, Methods M.3.3).

**Fig. 3.**
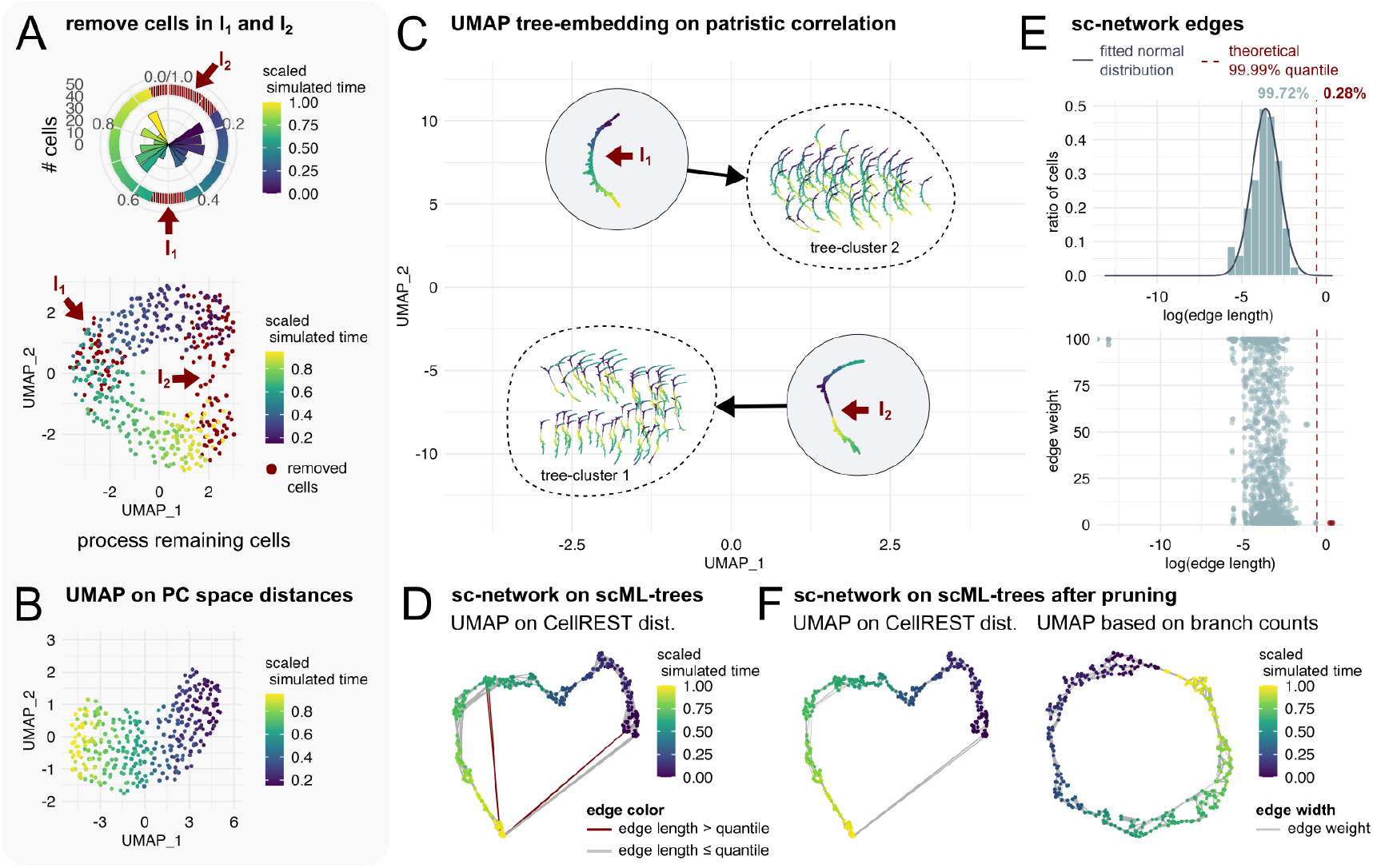
CellREST recovers non-tree-like dynamics under incomplete sampling. **A** The simulated circular differentiation process is sub-sampled by removing cells from two time intervals to mimic missing transient states. **B** Standard UMAP embedding of the remaining cells suggests an almost linear arrangement of cells. **C** UMAP tree-embedding of 100 scML-trees based on patristic-correlation; representative tree topologies highlight coverage of different trajectory segments. **D** UMAP visualizations of sc-network based on CellREST distances, with outlier edges (exceeding the quantile threshold) shown in red. **E** Histogram of logarithmized edge lengths in the sc-network with fitted normal distribution (top) and 99.99% quantile marked in red; edge weight vs length plot (bottom). **F** UMAP visualizations of the pruned sc-network based on CellREST distances (left) and shortest path distances using minimal branch counts between cells across nn-tree-graphs (right), with edge widths scaled by edge weight.

The standard UMAP embedding of the remaining cells failed to preserve the original circular topology, indicating its susceptibility in capturing the global structure in case of incomplete data (Fig. 3B).

Applying CellREST to the remaining cells, we again inferred 100 scML-tree topologies and identified two distinct tree-clusters. Each cluster effectively bridged one of the artificially introduced gaps (Fig. 3C, Fig. S3A). For the larger gap *I*_2_, trees attained noticeably longer internal branches to connect distant cell groups. As a result, the gap remained visible in the UMAP embedding based on CellREST distances. Nevertheless, the sc-network correctly reconnected the circular structure, unlike SNN-based graphs, which failed to capture the underlying topology (Fig. 3D, Fig. S3B). Notably, four logarithmized edge lengths in the sc-network exceeded the 99.99% quantile of the fitted normal distribution (Fig. 3E, Methods M.2.4). Crucially, removing these outlier edges preserved the gap-spanning connections and left CellREST distances unchanged, thereby leading to the same UMAP embedding (Fig. 3F). CellREST distances again achieved high Kendall’s tau correlations in comparison with simulation time (Fig. S3C).

The analyses of the simulated circular process and its subset demonstrate that scML-trees successfully approximate different parts of the underlying transcriptomic landscape (Fig. S3D). Together, these trees enable the reconstruction of trajectories that are non-tree-like or incompletely sampled.

In datasets containing unrelated cell populations (Fig. S4, Methods M.3.4), CellREST may generate connecting edges, as it does not enforce separation by design. However, such edges are typically among the longest in the sc-network, and hence can be identified as their lengths exceed the 99.99% quantile of the fitted normal distribution, as we show in a corresponding simulation scenario (Fig. S4D). These edges also tend to have low weights, indicating limited support across the nn-tree-graphs. This characteristic provides a practical strategy for filtering or down-weighting unreliable connections in downstream analyses.

### Experimental scRNA-seq Data

Simulated scRNA-seq data cannot fully capture the complexity of technical noise and biological variability present in experimental datasets (36). To complement our simulation-based evaluation, we applied CellREST to two benchmark datasets – mouse pancreatic endocrinogenesis (37) and peripheral blood mononuclear cells (PBMCs) (38) – as well as a renal cell carcinoma scRNA-seq data (39). To ensure comparability, each dataset was processed using the ‘standard’ Seurat and our CellREST workflow (Methods M.1).

#### Developing Mouse Pancreas

The mouse endocrinogenesis dataset (37) has become a benchmark for trajectory inference methods, as it is continuously sampled along the differentiation process from endocrine progenitors (EPs) to mature endocrine cell types Alpha, Beta, Delta, and Epsilon.

For the 3,696-cell dataset, we performed standard preprocessing and the resulting UMAP reflected the expected structure of the differentiation process (Fig. 4A, Methods M.4.1). Applying CellREST using 3,000 highly variable genes, we inferred 100 scML-trees, which revealed continuous but structured variation in a PCo tree-embedding (Fig. 4B). One group of trees displayed caterpillar-like topologies, reflecting a linear transition from early progenitors to terminal states (tree-cluster 3). Other trees captured alternative branching patterns, linking distinct EP subgroups to mature cell types via longer internal branches (tree-clusters 1,2 and 4). Tip instability analyses per cluster indicate that these groups represent different cutting locations in a potentially network-like transcriptomic landscape (Fig. S5A).

**Fig. 4.**
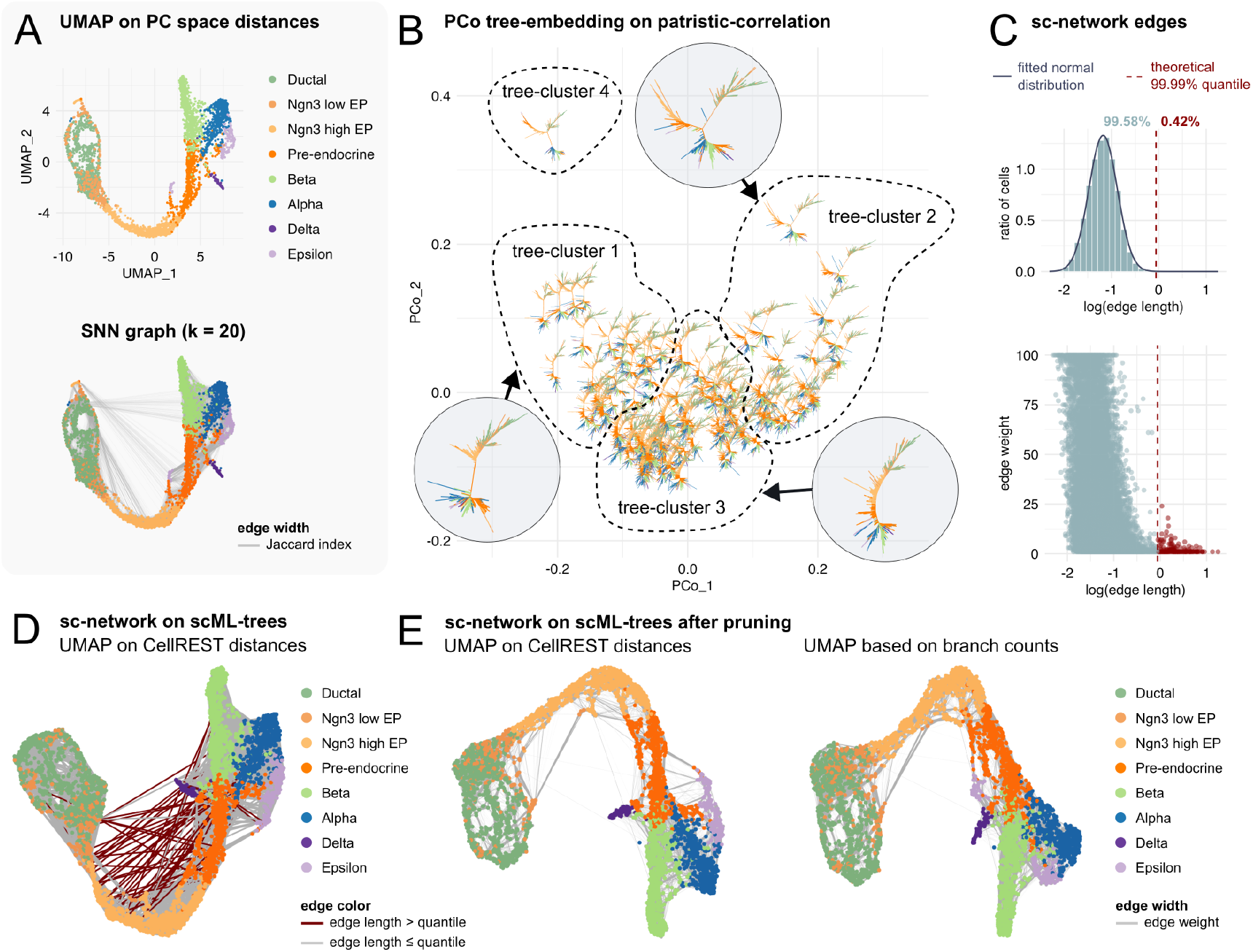
CellREST analysis of mouse endocrinogenesis scRNA-seq data. Abbreviations: EP = endocrine progenitor. **A** Standard UMAP embedding (top) and corresponding SNN graph edges (bottom), computed on PC space distances. Cells are colored by cell type annotation, edge widths reflect the Jaccard index between *k* neighborhood sets of connected cells. (37). **B** Principal Coordinates (PCo) tree-embedding of 100 scML-trees based on patristic-correlation, with representative tree visualized. **C** Histogram of logarithmized edge lengths in the sc-network (top), with a fitted normal distribution and the 99.99% quantile marked; edge weight vs length plot (bottom). **D** UMAP visualization of the full sc-network based on CellREST distances, with outlier edges (exceeding the quantile threshold) shown in red. **E** UMAP visualizations of the pruned sc-network based on CellREST distances (left) and shortest path distances using minimal branch counts between cells across nn-tree-graphs (right), with edge widths scaled by edge weight.

When combining all 100 scML-trees into a single sc-network, we identified a subset of outlier edges (0.42% of all edges), whose logarithmized lengths exceeded the 99.99% quantile of a fitted normal distribution (Fig. 4C). Following the strategy established in our simulations, we removed these edges and proceeded with a pruned version of the sc-network (Fig. 4D,E). Notably, some highly supported edges connecting cells outside the main differentiation wave remained (Fig. 4E) in the pruned sc-network and were also present in comparable SNN graphs (Fig. 4A, Fig. S5B). This suggests that the dataset may capture multiple, slightly variable differentiation waves converging to the same mature cell types. Visualizing the sc-network edges within the CellREST-informed UMAPs provides additional insight into sparsely sampled differentiation waves, particularly evident in the UMAP based on shortest path distances using minimal branch counts between cells across nn-tree-graphs (Fig. 4E).

#### Cell Type Clustering of Peripheral Blood Mononuclear Cells

We applied CellREST to a second benchmarking scRNA-seq dataset of peripheral blood mononuclear cells (PBMCs) (38) (Methods M.4.2), comprising distantly related cell types. In this dataset, cell types were experimentally identified using antibody-coated magnetic beads (Fig. S6A).

Following the evaluation strategy of Fu et al. (38), we assessed whether the CellREST-derived scML-trees (Fig. S6B) and the corresponding sc-network could be used to perform accurate cell type clustering. For this, we computed the Adjusted Rand Index (ARI) between experimentally identified cell types and computationally obtained cell clusters. Visual inspection of the sc-network before and after pruning of outlier edges (Fig. S6C) reveals a clear separation of experimentally defined cell types (Fig. S6D,E). Notably, several cell types retain dense internal connectivity within the pruned network. However, applying Louvain community detection (40) (Methods M.2.5) to the pruned sc-network with unweighted edges, produced slightly lower ARI scores compared to clustering on SNN graphs (Fig. S6F). This outcome might reflect the limitations of typically applied community detection methods (41), that do not take advantage of the distance information encoded by CellREST.

#### Putative Cells of Origin of Renal Cell Carcinoma

Finally, we applied CellREST to a renal cell carcinoma scRNA-seq dataset from Zhang et al. (39) (Methods M.4.3), which includes benign renal and tumor tissue from patients with chromophobe renal cell carcinoma (chRCC, 1 sample) and clear cell renal cell carcinoma (ccRCC, 7 samples). In this study, the authors aimed to identify putative cells of origin (P-COs) for each carcinoma type. Using gene expression profiles of epithelial cell types, they trained a random forest model to assign P-COs based on maximum prediction scores, proposing Proximal Tubule B (PT-B) cells as the P-COs for ccRCC and Intercalated (IC) cells for chRCC.

To focus the analysis, we sub-sampled the dataset to include only tumor cells (ccRCC and chRCC) and epithelial cell types. A standard UMAP embedding showed clear separation between epithelial cells and tumor cells, with the latter separating by patient origin (Fig. 5A).

**Fig. 5.**
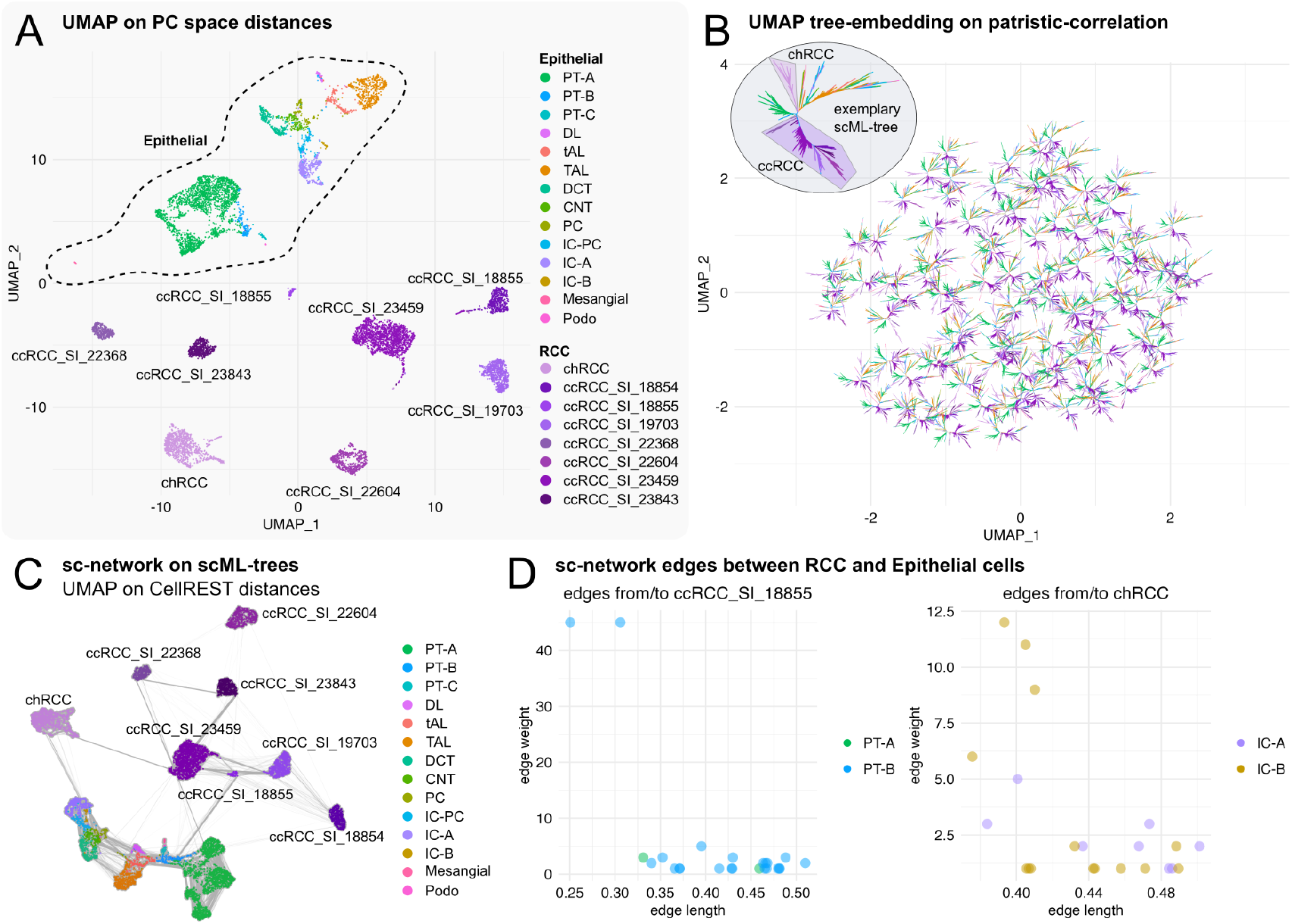
CellREST analysis of renal cell carcinoma data. Abbreviations: PT = proximal tubule, DL = descending limb, tAL = thin ascending limb, TAL = thick ascending limb, DCT = distal convoluted tubule, CNT = connecting duct, PC = principal cells, IC = intercalated cells, Podo = podocytes, RCC = renal cell carcinoma, cRCC = chromophobe RCC, ccRCC = clear cell RCC. **A** Standard UMAP embedding generated using Seurat, with cells are colored by original cell type annotations(39). ccRCC cells are further split by patient sample. **B** UMAP tree-embedding of 100 scML-trees based on patristic-correlation, with representative tree visualized. **C** UMAP visualizations of pruned sc-network based on CellREST distances, edge widths scaled by edge weights. **D** Edge length and weight of sc-network connections between epithelial cell types and ccRCC (left) or chRCC (right) cells from patient sample SI_18855.

Applying CellREST, we reconstructed 100 scML-trees, in which cells from the same RCC samples were often positioned within the same subtree. Additionally, chRCC cells consistently formed a distinct subtree, clearly separated from ccRCC cells (Fig. 5B). After pruning outlier edges (Fig. 5C, Fig. S7A–C) the resulting sc-network primarily connected epithelial cells among themselves and tumor cells within their subtype. Notably, we also observed direct connections between ccRCC cells and PT-B cells, including two prominent edges from patient samples SI_18855 and SI_18854 with high weights (weights 45 and weights 79, respectively) and moderate edge lengths (distances 0.251 and 0.306 and distances 0.198 and 0.223, respectively) (Fig. 5D) and lower weight edges from samples SI_19703, SI_22368 and SI_23459 (Fig. S7D). Likewise, chRCC cells showed direct but less strongly supported links to IC-A and IC-B cells (Fig. 5D). For two ccRCC samples (SI_22604, and SI_23843), no direct edges to epithelial types remained after pruning.

Using CellREST, we obtained a sc-network in which cells are connected if transitions between their gene expression profiles are considered likely based on phylogenetic inference. In the context of the carcinoma dataset, connections between healthy epithelial cells and tumor cells do not merely reflect transcriptomic similarity. Rather, they suggest that the gene expression profile of the linked epithelial cell could plausibly evolve into that of the connected carcinoma cell, providing a framework for identifying putative cells of origin.

## Discussion

With CellREST, we introduce a phylogenetic paradigm for analyzing single-cell transcriptomic data. CellREST implements a character-based phylogenetic framework that infers and analyzes gene expression state transitions between cells, thereby addressing a central challenge in trajectory inference – the lack of explicit ancestral information of a cell’s transcriptomic profile. Going beyond previous phylogenetic approaches (17, 18), CellREST utilizes and extends phylogenetic inference to address nontree-like transcriptomic landscapes by evaluating and integrating multiple similarly plausible single-cell-labeled Maximum Likelihood trees (scML-trees) into a unified singlecell network (sc-network). An implementation of CellREST is available as an R Package and integrates with the Seurat workflow.

Through simulation studies, we demonstrate that CellREST effectively recovers non-linear, circular trajectories – capabilities beyond many conventional trajectory or single-tree-based methods. In this context, CellREST distances between cells more accurately reflect the underlying temporal information than Euclidean distances computed from expression profiles, PC space, or UMAP embeddings. This improved resolution arises from their basis in model-inferred branch lengths and their definition as shortest paths, allowing them to follow curved transcriptomic trajectories. When applied to experimental data, such as mouse pancreatic endocrinogenesis, CellREST reveals converging differentiation waves and enhances their visualization via sc-network edges. Moreover, by identifying direct connections between cell types, CellREST can help to formulate hypotheses about putative progenitor cells or cells of origin, as shown in our analysis of renal cell carcinoma data.

The sc-network constructed by CellREST does not follow classical phylogenetic network approaches such as Neighbor Net (42, 43). This established method relies on predefined distances and often struggles to capture circular or noisy transcriptomic structures. Consensus Networks and other splits-based techniques (44) are especially sensitive to rogue taxa and noise, as demonstrated by the biologically uninformative Robinson-Foulds-based embeddings of scML-trees. Additionally, the scale of typical single-cell datasets exceeds the computational capabilities of many existing phylogenetic network implementations (45).

### Open questions and limitations

We are convinced that CellREST can be extended to other types of multi-model sequencing data through the use of phylogenetic partition models. Similarly, heterotachy models (46), which are designed to accommodate rate variation across lineages, may enhance the CellREST performance. However, the high dimensionality of the pseudo-alignments already pushes the limits of existing phylogenetic models and algorithms, and increasing model complexity risks introducing instability into tree inference. Beyond relying on stochastic variation between repeated maximum likelihood tree searches, future directions could include alternative strategies for exploring the scML-tree space, such as identifying terraces of quasi-likely trees (47) or employing Bayesian frameworks (48–50).

Currently, CellREST constructs pseudo-alignments by binning log-normalized gene expression values across a limited number of highly variable genes, that can be smaller than the number of considered cells. While variability is crucial for cell type discrimination, we suspect that including moderately or even uniformly expressed genes could benefit phylogenetic signal and contribute to a stable tree inference. Optimizing gene selection, binning strategies, and alternative normalization schemes will be important directions for improving the utility of the resulting pseudoalignments.

The connectivity of the resulting sc-network is determined by the user-defined parameter *B*, comparable to the role of *k* in the construction of *k*-nearest neighbor (NN) and corresponding shared-nearest neighbor (SNN) graphs. However, while *k* is often chosen without clear interpretability, *B* is more directly grounded in tree topology. Additionally, the current strategy for pruning outlier edges – based on the distribution of logarithmized edge lengths – would benefit from further statistical validation or refinement, such as using alternative criteria informed by phylogenetic saturation.

Our initial applications, namely CellREST-based visualizations and cell type clustering, represent only a starting point for a refined scRNA-seq analysis. While we value the ongoing developments in the field, such as improved normalization strategies and embedding techniques, we deliberately chose not to benchmark CellREST across diverse preprocessing pipelines. Instead, we focused on a standard workflow to emphasize that CellREST represents a *conceptual shift* in single-cell analysis. Moreover, the lack of appropriate experimental validation datasets currently limits comprehensive benchmarking. We therefore invite the scientific community to explore and build upon CellREST, particularly by utilizing its multi-layered information structure to rethink established analysis routines. At the same time, we acknowledge that the computational demands of repeated large-scale tree inference remain a limitation. However, with ongoing advances in scalable phylogenetic methods (51), we expect that CellREST will become increasingly accessible in practice. By fostering collaboration between the single-cell and phylogenetics communities, we hope to inspire the development of new tools that more accurately reflect the complexity of cell state transitions.

## Acknowledgments

Work in A.v.H.’s laboratory is supported by the Austrian Science Fund (Special Research Program SFB-F78, F 7811-B) and the Research Platform SinCeReSt - Single cell regulation of stem cells. We thank all members of the A.v.H.’s laboratory for their feedback and especially want to thank Florian Pflug and Stephanie Naas for insightful discussions.

## Author contributions

Conceptualisation: J.N., C.E. and A.v.H; Methodology: J.N., C.E.; Software: J.N.; Formal analysis: J.N., C.E.; Visualisation: J.N.; Writing - original draft: J.N.; Writing - review and editing: J.N., C.E. and A.v.H.

## Data and Code availability

The CellREST R package will be accessible on GitHub (https://github.com/jn-goe/CellREST).

## Methods

### M.1. Pre-Processing and Downstream Analysis

After dataset dependent quality control (see Section M.3 and Section M.4), the raw count matrix of a scRNAseq dataset of *m* cells is cell-wise log-normalized (‘`LogNormalize()`’) and a dataset dependent number *g* of highly variable genes (‘`nfeatures = g`’) is identified (‘`FindVariableFeatures()`’) using R package ‘Seurat’ (version 5.1.0) (21). Read imputation is performed on the complete log-normalized count matrix calling ALRA (27) via function ‘`RunALRA()`’ of R package ‘SeuratWrapper’ (version 0.3.5). ALRA applies singular vector decomposition on the measured count matrix to approximate it via a low-rank matrix. By assuming, that in this low-rank matrix, biological zeros admit values symmetrically distributed around zero, negative values can be used to compute a threshold to assign close-to-zero values as biological zeros.

After these pre-processing steps, we obtain an imputed log-normalized count matrix of *m* cells, that we subset to the *g* highly variable genes. Rows of the resulting matrix 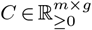 correspond to cells and columns to genes. This matrix serves as the starting point for the subsequent CellREST workflow.

For comparative reference, additional Seurat downstream analysis is performed, including gene-wise count scaling (‘`ScaleData()`’) and Principal Component Analysis (‘`RunPCA()`’), both performed on the selected highly variable genes. If not indicated otherwise, the top 50 principal components, spanning what we refer to the PC space, are used for dimensionality reduction via UMAP (22, 23).

### Nearest Neighbor Graphs

Principal components are also used to construct *k*-nearest neighbor (NN) and the corresponding shared-nearest neighbor (SNN) graphs (‘`FindNeighbors()`’). While the NN graph connects each cell to its *k*-nearest neighbors (by default *k* = 20, where the cell itself is counted as *k* = 1), the SNN only connects cells, as soon as they share a certain number of *k*-nearest neighbor. By default, only edges between cells are included in the SNN, as soon as the Jaccard index between the *k*-nearest neighbor sets is greater than 1*/*15. The Jaccard index is stored as edge attribute ‘weight’ in the resulting SNN. For the computation of shortest path distances on NN and SNN graphs, edges were additionally assigned an ‘edge length’ attribute, namely the distance between connected cells in the PC space, in which the graph was constructed. For clustering, by default, Louvain community detection is performed on SNN graph using function ‘`FindClusters()`’ for a specified resolution parameter ‘`res`’.

### M.2 CellREST Workflow

#### M.2.1. Transformation into a Phylogenetic Problem

For a scRNA-seq dataset with *m* cells, the log-normalized count matrix 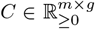 is subset to *g* highly variable genes and reformulated in phylogenetic terms as follows: For each gene *j*, expression levels across cells *c*_·*j*_ ∈ ℝ^*m*^ are uniformly binned into four discrete expression states (no, low, medium, and high expression) and subsequently renamed into pseudo-nucleotides (T,G,C, and A) by the following scheme:

| expression levels | expression state | pseudo-nucleotide |
| --- | --- | --- |
| $c_{ij} = 0$ | no expression | T |
| $c_{ij} \in \left(0, Q\left(\frac{1}{3}\right)\right]$ | low expression | G |
| $c_{ij} \in \left(Q\left(\frac{1}{3}\right), Q\left(\frac{2}{3}\right)\right]$ | medium expression | C |
| $c_{ij} \in \left(Q\left(\frac{2}{3}\right), Q(1)\right]$ | high expression | A |

for a cell *i*, where *Q*(*p*) = *p* · (max_*i*_(*c*_*ij*_) − min_*i*_(*c*_*ij*_)) + min_*i*_(*c*_*ij*_) is the theoretical *p*-quantile of the uniform distribution *U* [min_*i*_(*c*_*ij*_), max_*i*_(*c*_*ij*_)]

The resulting pseudo-alignment, which comprises *m* pseudo-taxa (= cells = rows) and *g* pseudo-sites (= genes = columns), is stored as a FASTA file using R package ‘seqinr’ (version 4.2-36) (52) and can be used for subsequent phylogenetic inference.

#### M.2.2. Phylogenetic Inference

Preliminary evaluations on test datasets indicated that the Maximum Likelihood (ML) method is best suited for the evaluation of generated pseudo-alignments and identified the nucleotide substitution model GTR+F+G4 as the best-fit using ModelFinder (53), as implemented in IQ-TREE2 (24) (version 2.2.2.4). GTR (General Time Reversible) is a flexible model that allows different substitution rates between all pairs of nucleotides, F (empirical base frequencies) incorporates observed nucleotide frequencies from the data, and G4 (Gamma distribution with four rate categories) accounts for rate variation across sites. To maintain consistency and enable comparability across results, this evolutionary model class is fixed for all analyses in this manuscript.

For each pseudo-alignment, *n*_*trees*_ = 100 single-cell-labeled ML-trees (scML-trees) are inferred using IQ-TREE2 (24) (version 2.2.2.4). This can be done by calling the IQ-TREE option ‘`--runs NUM`’ or using the provided ‘run_iqtree.sh’ script in the CellREST package, which parallelizes the *n*_*trees*_ tree inference procedures. To enhance computational efficiency, we perform tree reconstruction using IQ-TREE’s ‘`-fast`’ option using a fix number of cores (‘`N_CORES`’) and a pre-defined seed (‘`SEED`’) for reproducibility:

iqtree2 -s FASTA_FILE -m GTR+F+G4 -nt N_CORES -seed SEED -fast

Reconstructed trees are processed and visualized using R packages ‘ape’ (version 5.8), ‘castor’ (version 1.8.2), ‘ggtree’ (version 3.12.0)(54) and tips are colored by data-dependent cell annotations.

#### M.2.3. Comparing and Integrating scML-trees

There are several methods for comparing phylogenetic trees, depending on the specific aspect of interest – such as topology, branch lengths, or leaf labeling. For instance, the Robinson-Foulds (RF) distance (here computed using R package ‘treespace’, version 1.1.4.3 (30)) measures structural differences between two trees by counting the number of splits (bipartitions) that are unique to each tree. While the RF distance quantifies purely structural differences, the below introduced patristic-correlation is a pseudo-distance that incorporates branch length information.

##### Patristic Correlation Tree Pseudo-Distance

Given one scML-tree, the evolutionary distance – here in the form of expression profile changes – between two different cells can be determined by calculating the sum of the branch lengths along the shortest path that connects them, i.e., the ‘patristic’ distance. By computing the patristic distances between all pairs *i* and *j* of *m* cells given a tree *T*, one obtains a matrix of cell-to-cell patristic distances

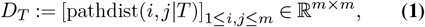

which can be vectorized into one high-dimensional patristic distance vector 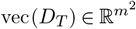.

Now, to compare two trees *T*_*i*_ and *T*_*j*_, that comprise the same set of *m* cells, we first calculate their corresponding patristic cell-distance vectors, denoted by 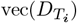 and 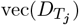. Following the approach of Chen et al. (28), we define a similarity measure based on the sample Pearson’s correlation coefficient between the two patristic celldistance vectors as

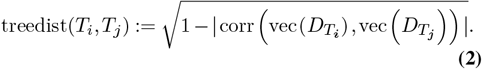

This is a pseudo-distance since there can exist two distinct trees *T*_*i*_ ≠ *T*_*j*_ for which treedist(*T*_*i*_, *T*_*j*_) = 0. By computing the patristic-correlation between all pairs of *n*_*trees*_ trees, one obtains a tree-distance-matrix

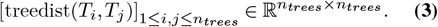

We decided to utilize a correlation measure, since it is more robust to outliers and noise compared to, e.g., the Euclidean distance computed between high-dimensional distance vectors.

##### Embedding and Hierarchical Clustering of scML-trees

A tree-distance-matrix computed via pairwise comparison of all scML-trees can be used to embed the trees into a lower dimension in order to visualize and cluster them. To do so, one can use Principal Coordinate Analysis (PCoA, also called Multidimensional Scaling), which is the equivalent of Principal Component Analysis for distances instead of coordinates, or UMAP, which can be applied on distances as well as on coordinates.

For the UMAP computation we use function ‘`umap()`’ of R package ‘uwot’ (version 0.2.3) setting parameters as they are the default in UMAP computations with R package ‘Seurat’, i.e., ‘`n_neighbors = 30`’, ‘`min_dist = 1`’, and ‘`seed = 42`’ for reproducibility.

For both PCoA and UMAP we use the first two components for visualization and hierarchical clustering of scML-trees with function ‘`hclust()`’ of R package ‘stats’ (version 4.4.0) gaining a data-specific number of clusters ‘`k`’ using ‘`cutree()`’.

##### Tip-instability Measure

To identify rogue taxa, that lead to very different patristic distances across a group of scML-trees, we compute the tip instability measure using function ‘`TipInstability()`’ in R package ‘Rogue’ (version 2.1.6) by Smith et al. (35).

#### M.2.4. Single-cell-network Construction

##### Construction of nn-tree-graphs

As a noise-robust representative of a scML-tree we suggest to transform a tree into a nn-tree-graph. To do so, for a tree *T* with *m* cells, we initiate a graph with corresponding *m* vertices. For each cell, we add an edge from the corresponding vertex to all vertices that are at most *B* branches apart. For all analyses of this study, we set *B* = 4. The resulting nn-tree-graph *G* is defined by its adjacency matrix *A*(*G*) ∈ {0, 1} ^*m×m*^, where an entry *A*(*G*)_*ij*_ = 1 implies an edge (*i, j*) connecting vertices *i* and *j*.

For each edge (*i, j*), we additionally store the patristic distance between the two connected cells *i* and *j* as *length* attribute

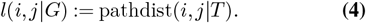

##### Union of nn-tree-graphs into a sc-network

For a group of *n*_*trees*_ trees 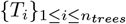 defined on the same set of cells *m*, we can construct the union across respective nn-tree-graphs 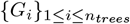 including the same set of vertices *m*. In more detail, we entry-wise sum all adjacency matrices of considered nn-tree-graphs yielding a new, weighted adjacency matrix

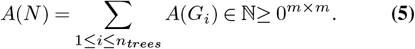

This defines the single-cell network (sc-network) *N*, where a positive value *A*(*N*)_*ij*_ *>* 0 again corresponds to an edges (*i, j*) connecting vertices *i* and *j*. The exact value *A*(*N*)_*ij*_ determines the **edge weight** of edge (*i, j*), i.e., the number of nn-tree-graphs, in which this edge was observed.

For each edge (*i, j*) of the sc-network *N*, we additionally compute a representative **edge length** as the minimum of edge lengths across all considered nn-tree-graphs, in which the edge (*i, j*) was observed,

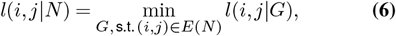

where *E*(*N*) denotes the set of all edges of graph *N*.

##### CellREST Distance

Finally, the edge lengths of a sc-network can be used to compute a shortest path distance between cells *i* and *j* in the sc-network *N*, which we will refer to as CellREST distance,

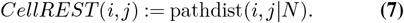

Computed between all pairs of *m* cells, we obtain the CellREST cell-distance-matrix

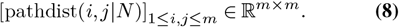

##### Fitting a Normal Distribution to Logarithmized Edge Lengths

To fit a theoretical distribution to the empirical distribution of edge lengths *l*^*N*^ in sc-network *N*, we first log-transform all edge lengths such that a symmetric normal distribution can be fitted. The median of the log-transformed edge lengths is used as an estimate for the mean of the normal distribution

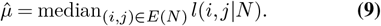

To ensure robustness against large outliers, we make use of the fact that, under the normal distribution, the in-terquantile range is proportional to the standard deviation.Therefore, we estimate the standard deviation as

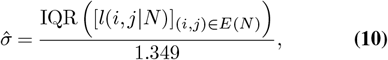

where [*l*(*i, j* | *N*)]_(*i,j*) ∈ *E*(*N*)_ is the vector containing all logarithmized edge lengths of network *N*, and IQR(*x*) := *Q*_*x*_(3*/*4) − *Q*_*x*_(1*/*3) the interquartile range of a vector *x* with *Q*_*x*_(1*/*4) and *Q*_*x*_(3*/*4) being the empirical lower and upper quartiles of *x*, respectively.

We perform pruning on edges, that have a length greater than the theoretical 99.99% quantile of the fitted normal distribution 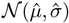.

#### M.2.5. Sc-network-based Downstream Analysis

##### Distance-based Embeddings

We use the CellREST distance-matrix for visualizations of cells via dimensionality reduction methods PCoA and UMAP (as described for the tree-distance-matrix in Section M.2.3). Using the coordinates of embedded cells, corresponding network edges can be plotted using R package ‘igraph’ (version 2.0.3) (55) to additionally visualize the connectivity of the sc-network. As an alternative visualization, we also include UMAP embeddings computed on shortest path distances after replacing the minimal patristic distance by the minimal branch count between cells as an edge length attribute in the sc-network.

##### Cell Clustering

To perform clustering on the CellREST sc-network, we transform the network to an unweighted adjacency matrix (entries either 0 or 1 depending on the existence of an edge) and store it as ‘Graph’ object in ‘obj@graphs$CellREST’ of the corresponding Seurat object ‘`obj`’. Subsequently, we perform Louvain clustering by calling Seurat function ‘`FindClusters()`’ specifying ‘`graph.name = ‘CellREST’`’.

### M.3. Simulation with Dyngen

For our simulation studies we use R package ‘dyngen’ (version 1.0.5) (33) to generate scRNA-seq data following predefined trajectories. Dyngen integrates a gene regulatory network and a module network that defines the ‘backbone’ topology, the main trajectory the simulated cells will follow. To generate realistic gene expression levels along a continuous simulation time scale, transcription, splicing and translation is simulated in a chain of reactions via Gillespie’s stochastic simulation algorithm (34). Finally, cells and mRNA molecules are sampled to imitate the technical procedure of sequencing.

For the CellREST binning procedure 300 highly variable genes are used for all simulated datasets.

#### M.3.1. Simple Circular Process

Initially, 2,000 cells are simulated based on the ‘cycle_simple’ backbone, the simulation model was initialized using function ‘`initialise_model()`’ setting ‘`num_tfs = 500`’ (number of transcription factors), ‘`num_targets = 500`’ (number of target genes) and ‘`num_hks = 1000`’ (number of housekeeping genes) with seed 14 and running 1 simulation with census interval 0.1, i.e., ‘`simulation_default(experiment_params = simulation_type_wild_type(num_simulations` `= 1), census_interval = .1)`’. Since one simulation approximately covers around 2.5 repetitions of the circle backbone, cells are subset to only one cycle loop and sub-sampled to 150 cell from each of the three milestone states *s*_1_, *s*_2_ and *s*_3_. Finally, the simulation time on the remaining 450 cells is scaled to [0, 1] and Seurat downstream analysis is performed as described above.

#### M.3.2. Correlation of Simulation Time Differences in Circular Processes

To evaluate how well CellREST distances, Euclidean distances in embeddings and shortest path distances on NN or SNN graphs (construction see Section M.1) encode the simulation time, we performed a correlation based evaluation.

First we start by constructing a matrix of cycle-aware simulation time differences between all *m* cells. To do so, we compute differences in simulation time between all pairs of cells and store them in matrix *D*^*forward*^ ∈ ℝ^*m×m*^. To take into account the circular shape of the topology, i.e., the fact that simulation time 1 and 0 coincide, we also compute the simulation time differences on the reversed simulationtime *D*^*backward*^ = −*D*^*forward*^%1, using modulo operator %. The final simulation time difference matrix *D*^*time*^ is then defined as entry-wise minimum

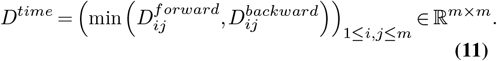

To compare *D*^*time*^ with other method-specific distance-matrices *D*^*method*^, for each cell *i* we compute the Kendall’s correlation coefficient *τ* between corresponding rows of *D*^*time*^ and *D*^*method*^

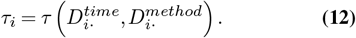

Across all *m* cells this yields a set of Kendall’s correlation coefficients {*τ*_*i*_} _1*≤i≤m*_ that can be visualized via boxplots using ‘ggplot2’ (version 3.5.1).

#### M.3.3. Discontinuous Circular Process

To simulated missing ancestral information, simulated cells from a circular process are subset by deleting those cells of scaled simulation time in intervals *I*_1_ = (0.45, 0.5) and *I*_2_ = (0.15, 0.95). Downstream analysis is repeated on the remaining 314 cells.

#### M.3.4. Disconnected Cell Groups

500 cells are simulated based on the ‘disconnected’ backbone, the simulation model was initialized setting ‘`num_tfs = 500`’, `num_targets = 500`’ and `num_hks = 1000`’ with seed 14 and running 10 simulations, i.e., ‘`simulation_params = simulation_default(experiment_params = simulation_type_wild_type(num_simulations` `= 10))`’. Running several simulations mimics the biological scenarios that multiple cell lineages follow and are sampled along the same trajectory.

### M.4. Experimental Datasets

For all experimental datasets 3,000 highly variable genes are used for Seurat as well as CellREST analyses. For the visualization of trees, we additionally used function ‘`groupOTU()`’ of package ‘ggtree.’

#### M.4.1. Mouse Pancreatic Endocrinogenesis

The ‘anndata’ object including pre-processed scRNA-seq counts and cell type annotation of the developing mouse pancreas sampled from embryonic day 15.5 published by Bastidas-Ponce et al. (37) was imported using Python package ‘scvelo’ (version 0.3.3) (56) via function ‘`scvelo.datasets.pancreas()`’ and loaded into R via package ‘anndata’ (version 0.7.5.6). A Seurat object is initialized importing raw counts and metadata, comprising 3,696 cells.

#### M.4.2. Benchmark PBMC cells

Clustering benchmark PBMC data from Fu et al. (38) was downloaded from Zenodo (57). A Seurat object is initialized using raw counts imported from ‘Liu_raw_data.csv’ and cell type annotation imported from ‘Liu_purified_celltype.csv’. Only genes that were measured in at least one cell are retained yielding a dataset comprising 9,266 cells. For the comparison of cell type annotations by Louvain community detection obtained clusterings the Adjusted Rand Index (ARI) is computed using R package `mclust` (version 6.1.1).

#### M.4.3. Human Renal Carcinoma Cancer

Data of clear and chromophobe renal cell carcinoma (RCC) samples and benign tissue published by Zhang et al. (39) were downloaded from the National Center for Biotechnology Information Gene Expression Omnibus (GEO) under study number GSE159115. Per sample, genes are filtered following the processing described in Zhang et al. (39), i.e., only genes detected in at least five cells are retained and uninformative mitochondrial, ribosomal, and sex genes are removed. Subsequently, only high-quality cells listed in the provided metadata ‘GSE159115_ccRCC_anno.csv’, ‘GSE159115_chRCC_anno.csv’ and ‘GSE159115_normal_anno.csv’ are kept for the subsequent analysis. All samples are merged (without batch-correction) into one Seurat object comprising 29,474 cells. Prior to Seurat downstream analysis and application of CellREST, the dataset is subset to 5,000 randomly sampled cells, which are assigned as tumor cells, and all 3,956 cells that are assigned as Epithelial cells across all samples.

## Supplementary Figures

**Fig. S1.**
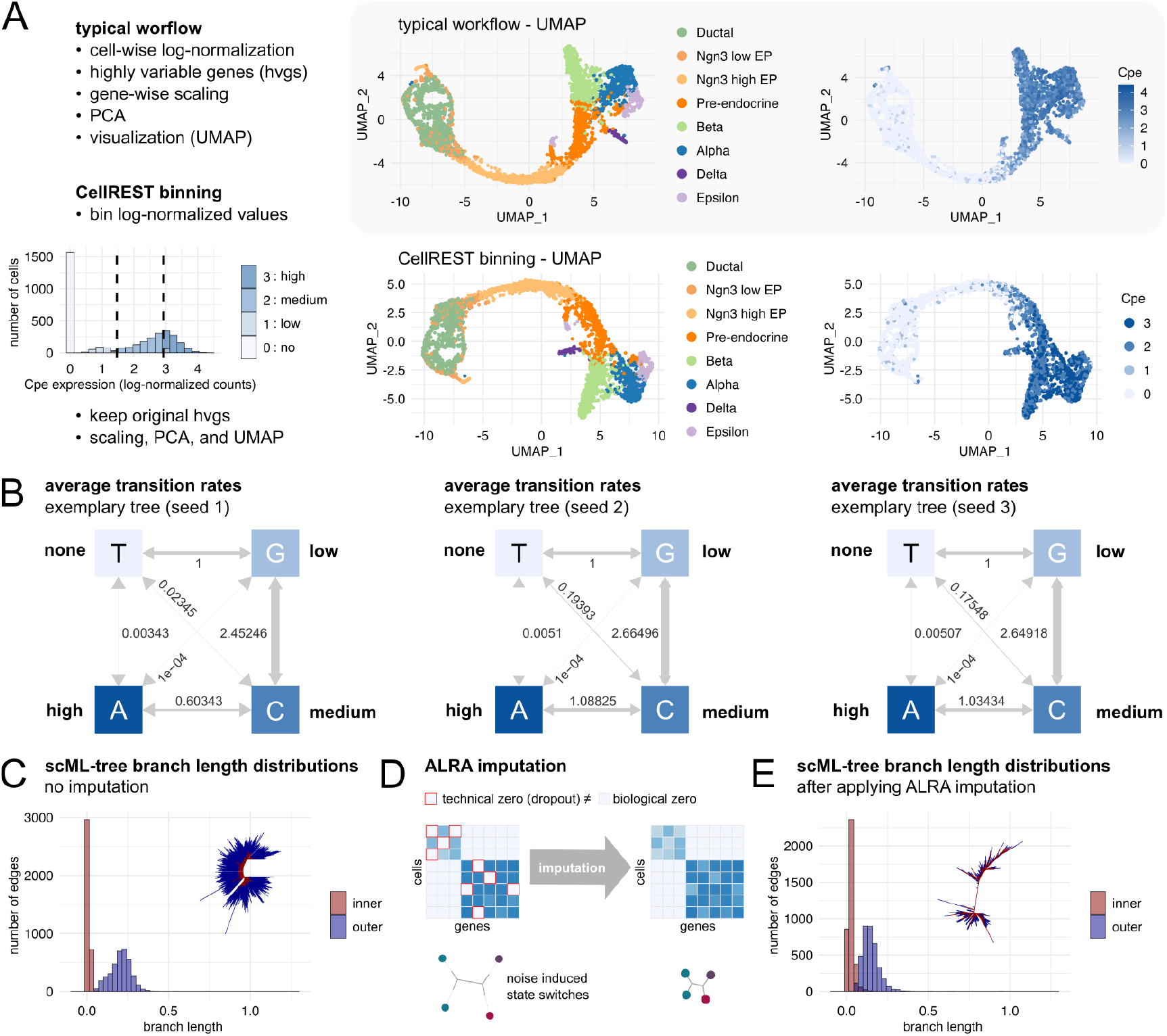
Details on CellREST workflow. Abbreviations: EP= endocrine progenitor. **A** Expression values per genes binned into four categories and mapped to numerical values 0,1,2,3 result in comparable UMAP to those based on log-normalized expression profiles. Exemplary UMAPs with cell type annotations (middle) and hormone processing marker gene Cpe (right) expression of endocrinogenesis scRNA-seq data from (37). **B** Average transition rates of inferred GTR models for an exemplary inferred scML-trees. **C** The distribution of branch lengths of scML-trees computed on log-normalized counts. **D** Sketch of ALRA imputation (27) and its effect on branch lengths in scML-trees. **E** scML-tree branch lengths decreased after ALRA imputation.

**Fig. S2.**
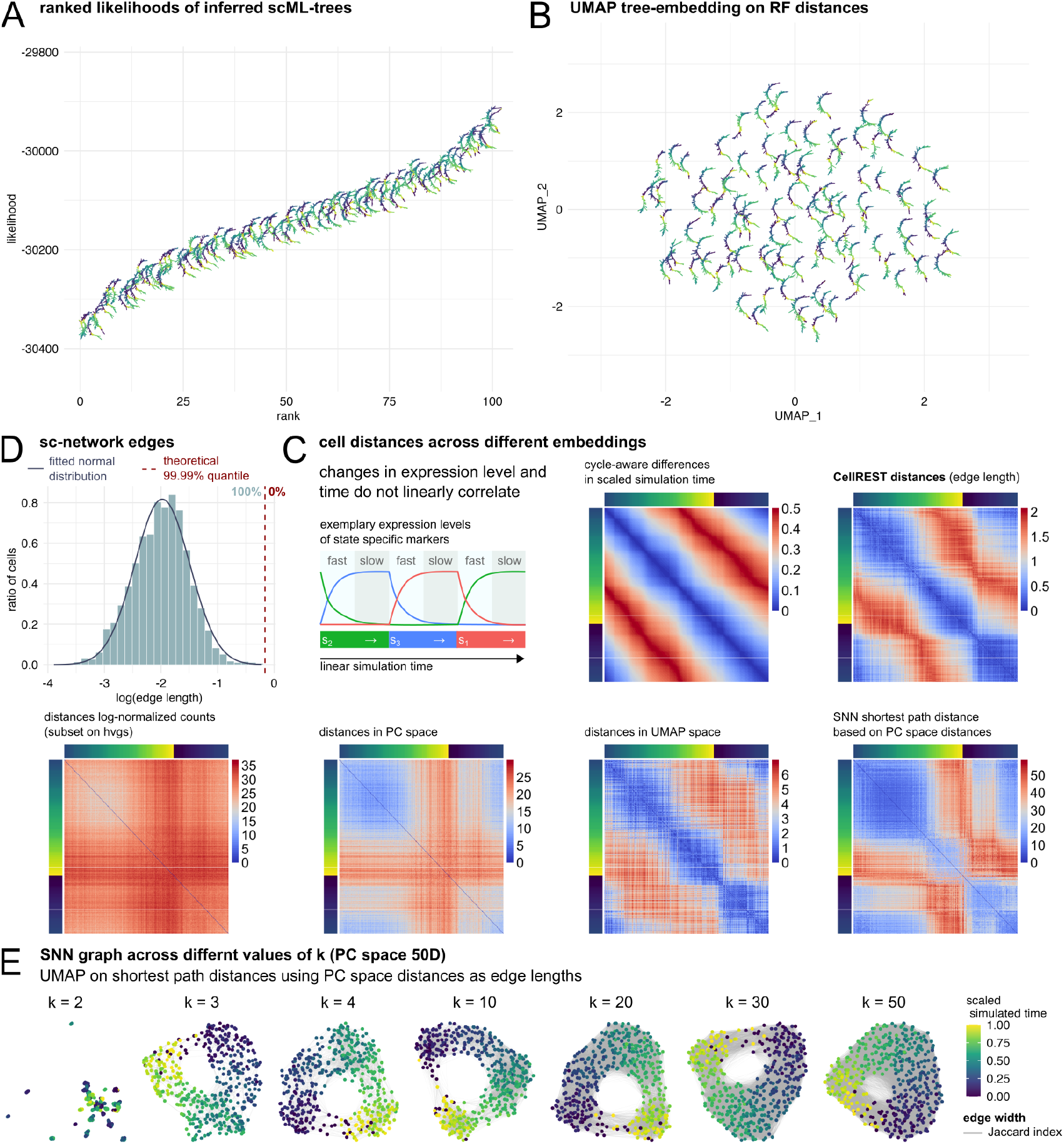
Additional analyses on the simulated circular process. **A** scML-trees ranked by likelihood. **B** UMAP tree-embedding based on Robinson Foulds distances. **C** Cell-to-cell distance heatmaps based on differences in simulation time (cycle-aware) and method-specific distances. **D** Histogram of logarithmized edge lengths of sc-network with fitted normal distribution and 99.99% quantile marked in red. **E** SNNs constructed across a range of *k* embedded using PC space distances, where edge widths are scaled by the Jaccard index between the *k* neighborhood sets of connected cells.

**Fig. S3.**
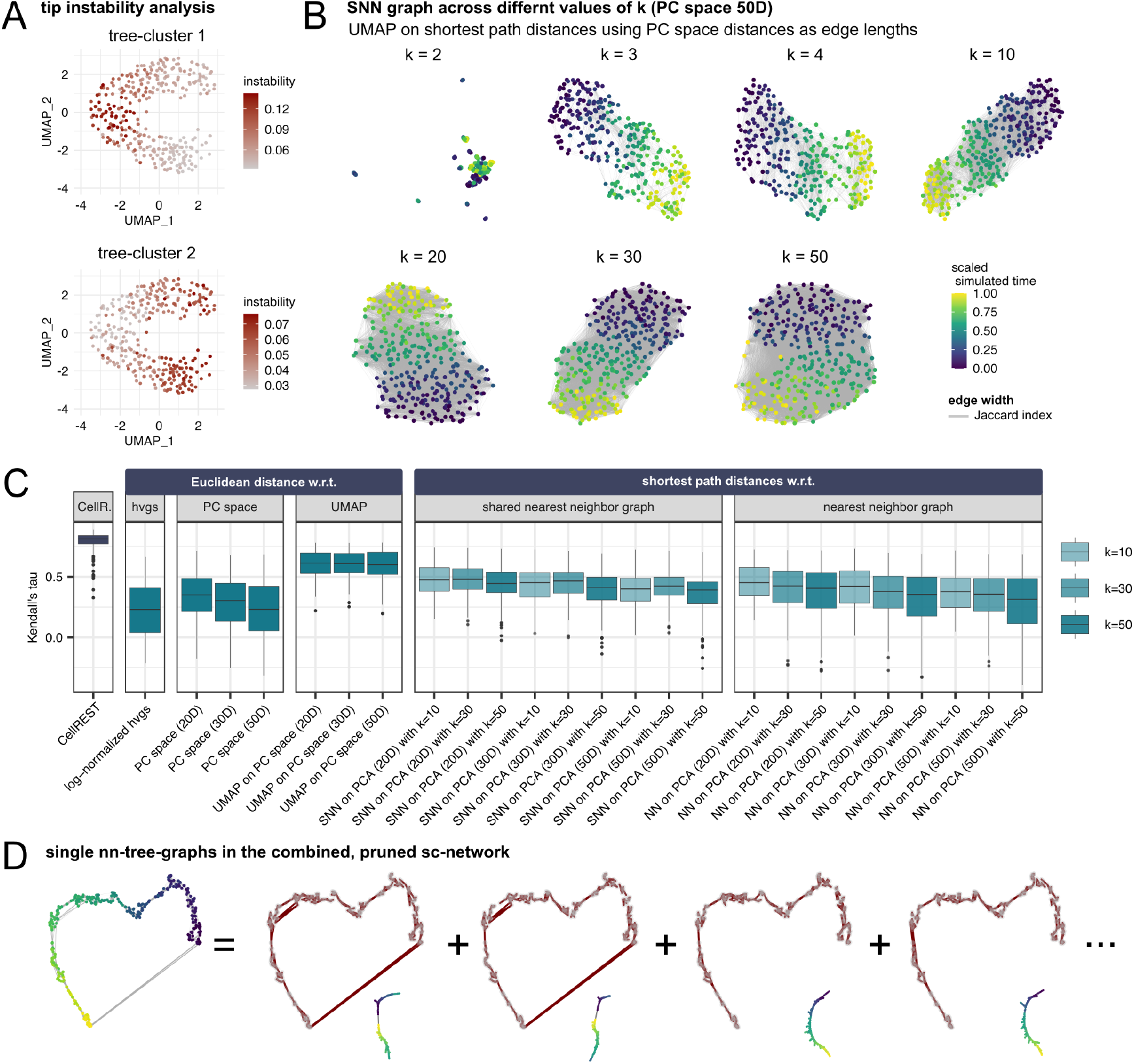
Additional analyses on simulated subset circular process. **A** Tip instability analysis for tree-clusters from Fig. 3C. **B** SNNs constructed across a range of *k* embedded using PC space distances, where edge widths are scaled by the Jaccard index between the *k* neighborhood sets of connected cells. **C** Kendall’s tau correlation between cell orderings based on differences in simulation-time (cycle-aware) and embedding-specific distances. **D** Exemplary tree-graphs approximate different parts of the trajectory network combined in the CellREST sc-network.

**Fig. S4.**
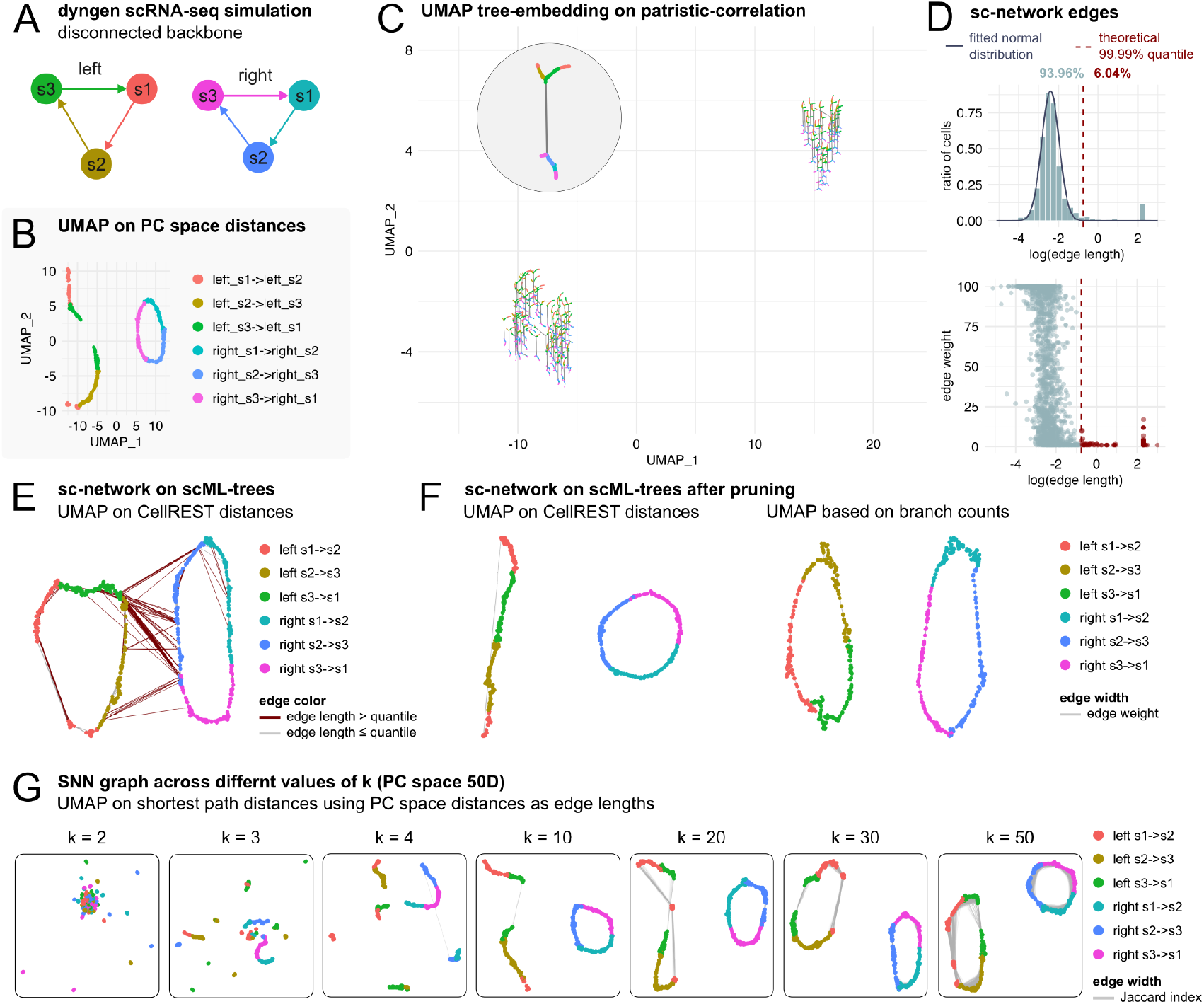
Additional analyses on simulated disconnected processes. **A** Setup for dyngen simulation of two unrelated circular processes. **B** UMAP computed on PC space distances following a typical downstream analysis. **C** UMAP tree-embedding of 100 scML-trees using patristic-correlation with detail on exemplary scML-tree. **D** Histogram of logarithmized edge lengths in the sc-network with fitted normal distribution (top) and 99.99% quantile marked in red; edge weight vs length plot (bottom). **E** UMAP visualizations of sc-network based on CellREST distances, where edges are red if their length exceeds the fitted quantile threshold. **F** UMAP visualizations of pruned sc-network based on CellREST distances (left) and shortest path distances using minimal branch counts between cells across nn-tree-graphs (right), where edge widths are scaled by the edge weight attribute. **G** SNN constructed across a range of *k* embedded using PC space distances, where edge widths are scaled by the Jaccard index between the *k* neighborhood sets of connected cells.

**Fig. S5.**
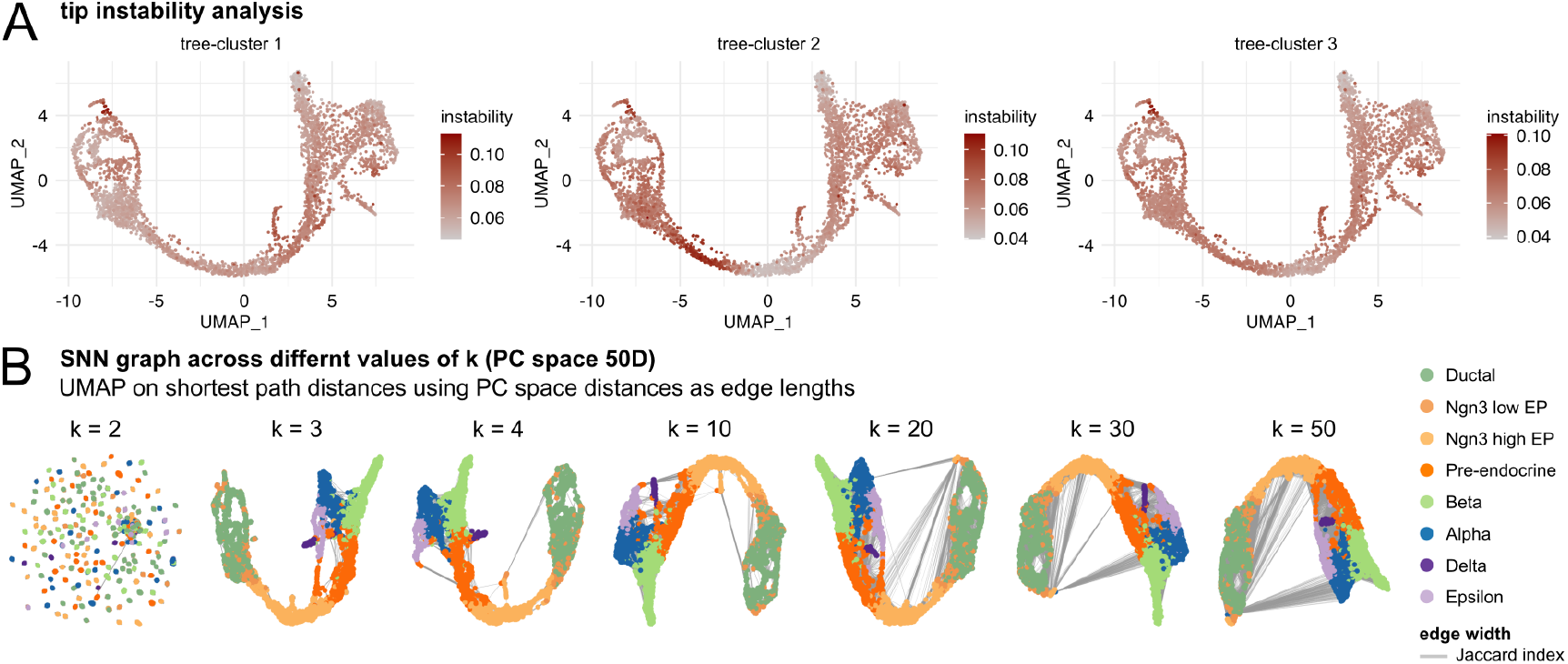
Additional analyses on mouse endocrinogenesis data. Abbreviations: EP= endocrine progenitor. **A** Tip instability analysis across tree-clusters as shown in Fig. 4C. **B** SNNs constructed across a range of *k* embedded using PC space distances, where edge widths are scaled by the Jaccard index between the *k* neighborhood sets of connected cells.

**Fig. S6.**
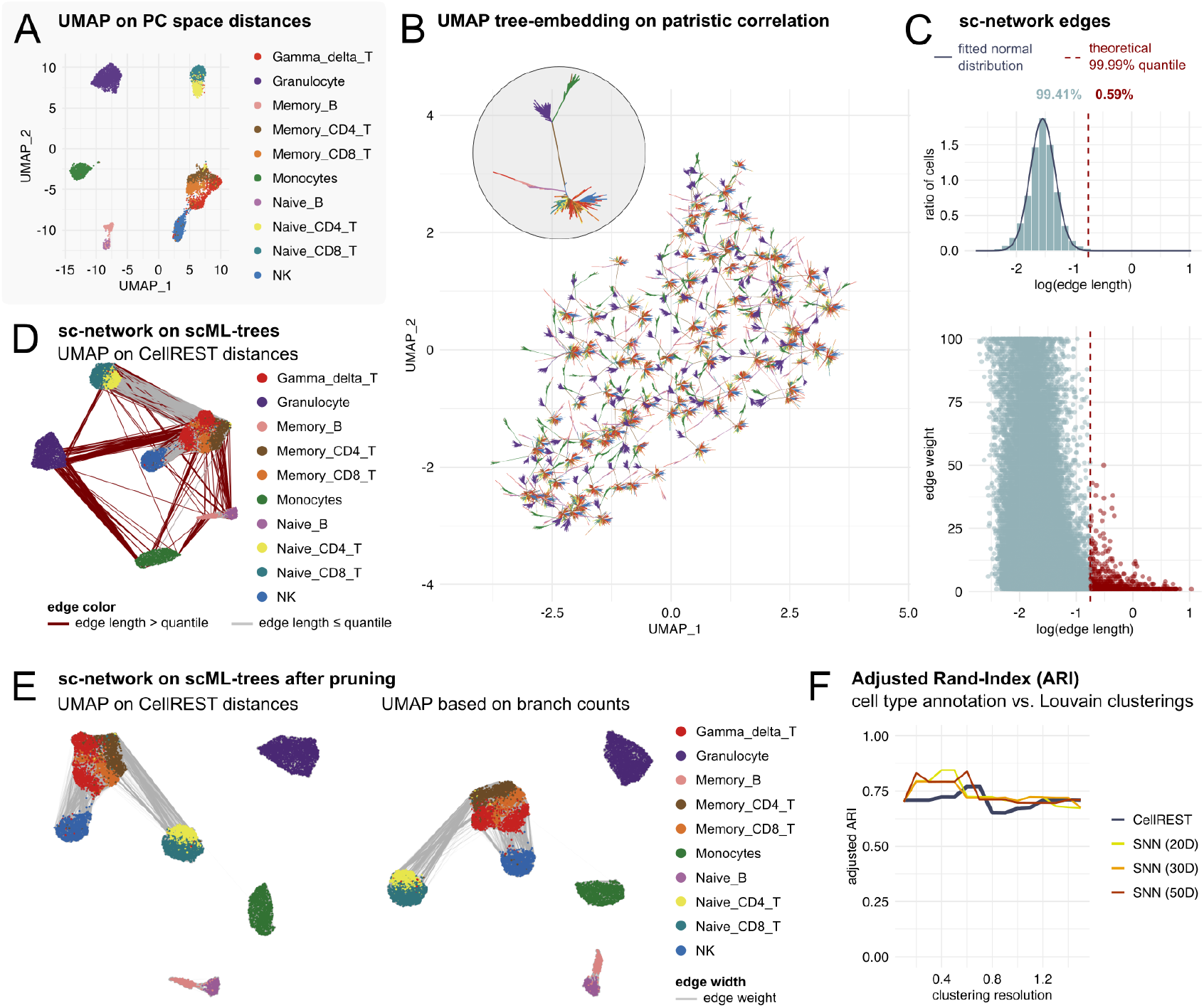
Additional analyses on blood cell data. **A** UMAP obtained by typical downstream analysis using Seurat color-coded by cell type annotation (38). **B** Tree-embedding of 100 scML-trees using patristic-correlation. **C** Histogram of logarithmized edge lengths in the sc-network with fitted normal distribution (top) and 99.99% quantile marked in red; edge weight vs length plot (bottom). **D** UMAP visualizations of sc-network based on CellREST distances, where edges are red if their length exceeds the fitted quantile threshold. **E** UMAP visualizations of pruned sc-network based on CellREST distances (left) and shortest path distances using minimal branch counts between cells across nn-tree-graphs (right), where edge widths are scaled by the edge weight attribute. **F** Adjusted rand index analysis for Louvain clusterings on SNN graphs and CellREST sc-network across a range of clustering resolutions.

**Fig. S7.**
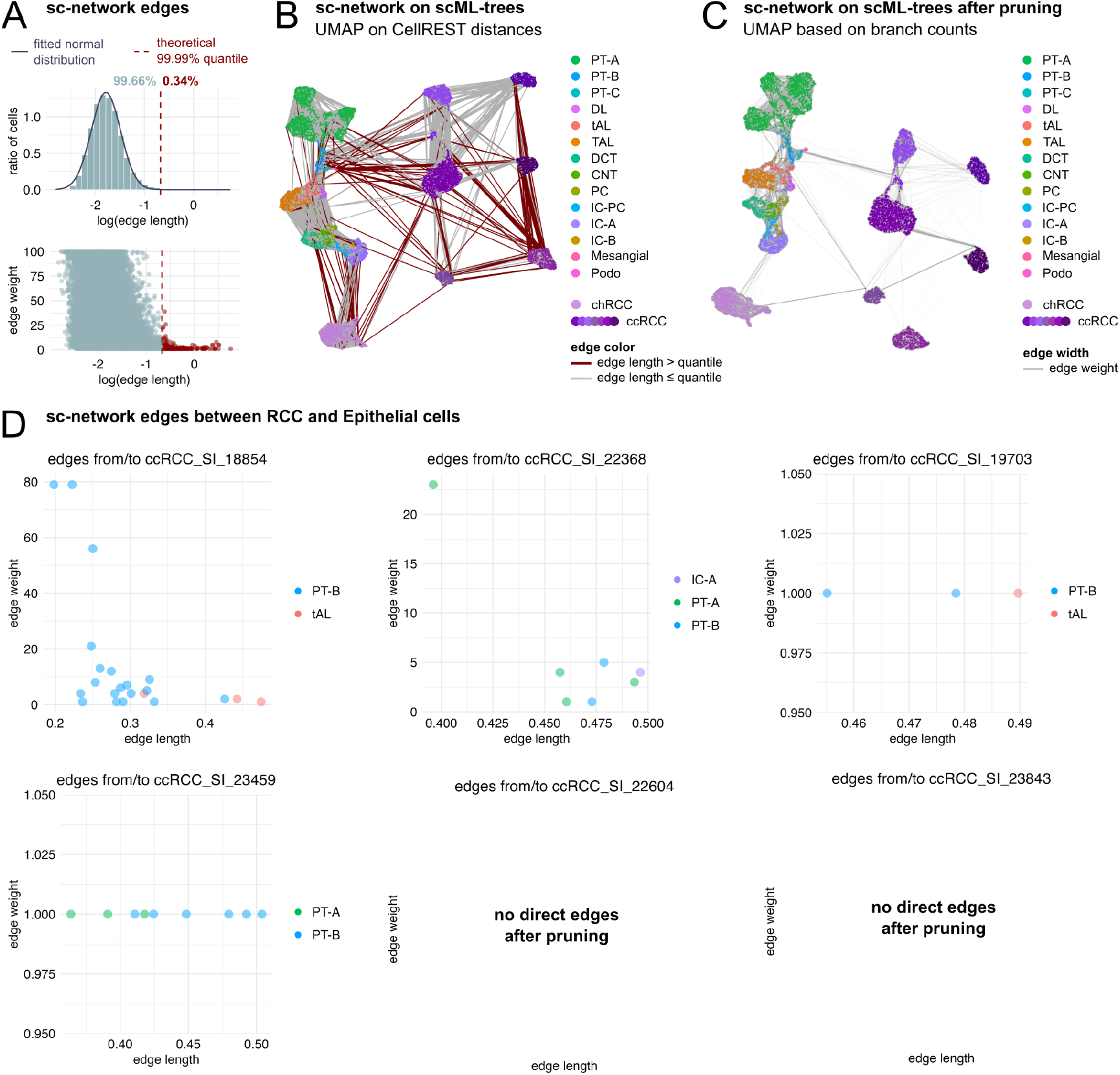
Additional analyses on renal cell carcinoma. Abbreviations: PT = proximal tubule, DL = descending limb, tAL = thin ascending limb, TAL = thick ascending limb, DCT = distal convoluted tubule, CNT = connecting duct, PC = principal cells, IC = intercalated cells, Podo = podocytes, RCC= renal cell carcinoma, cRCC = chromophobe RCC, ccRCC = clear cell RCC. **A** Histogram of logarithmized edge lengths in the sc-network with fitted normal distribution (top) and 99.99% quantile marked in red; edge weight vs length plot (bottom). **B** UMAP visualizations of sc-network based on CellREST distances, where edges are red if their length exceeds the fitted quantile threshold. **C** UMAP visualization of pruned sc-network based on shortest path distances using minimal branch counts between cells across nn-tree-graphs, where edge widths are scaled by the edge weight attribute. **D** Length and weight of sc-network edges that connect ccRCC cells to epithelial cell type clusters.

## References

1. Trapnell, C. et al. The dynamics and regulators of cell fate decisions are revealed by pseudotemporal ordering of single cells. Nature Biotechnology 32, 381–386 (2014). DOI: 10.1038/nbt.2859.

2. Qiu, X. et al. Reversed graph embedding resolves complex single-cell trajectories. Nature Methods 14, 979–982 (2017). DOI: 10.1038/nmeth.4402.

3. Cao, J. et al. The single-cell transcriptional landscape of mammalian organogenesis. Nature 566, 496–502 (2019). DOI: 10.1038/s41586-019-0969-x.

4. Street, K. et al. Slingshot: cell lineage and pseudotime inference for single-cell transcriptomics. BMC Genomics 19, 477 (2018). DOI: 10.1186/s12864-018-4772-0.

5. Setty, M. et al. Characterization of cell fate probabilities in single-cell data with Palantir. Nature Biotechnology 37, 451–460 (2019). DOI: 10.1038/s41587-019-0068-4.

6. Chen, H. et al. Single-cell trajectories reconstruction, exploration and mapping of omics data with STREAM. Nature Communications 10, 1903 (2019). DOI: 10.1038/s41467-019-09670-4.

7. Lange, M. et al. CellRank for directed single-cell fate mapping. Nature Methods 19, 159–170 (2022). DOI: 10.1038/s41592-021-01346-6.

8. Weiler, P., Lange, M., Klein, M., Pe’er, D. & Theis, F. CellRank 2: unified fate mapping in multiview single-cell data. Nature Methods 21, 1196–1205 (2024). DOI: 10.1038/s41592-024-02303-9.

9. Faure, L., Soldatov, R., Kharchenko, P. V. & Adameyko, I. scFates: a scalable python package for advanced pseudotime and bifurcation analysis from single-cell data. Bioinformatics 39, btac746 (2023). DOI: 10.1093/bioinformatics/btac746.

10. Saelens, W., Cannoodt, R., Todorov, H. & Saeys, Y. A comparison of single-cell trajectory inference methods. Nature Biotechnology 37, 547–554 (2019). DOI: 10.1038/s41587-019-0071-9.

11. Saitou, N. & Nei, M. The neighbor-joining method: a new method for reconstructing phylogenetic trees. Molecular Biology and Evolution 4, 406–425 (1987). DOI: 10.1093/oxfordjournals.molbev.a040454.

12. Gascuel, O. BIONJ: an improved version of the NJ algorithm based on a simple model of sequence data. Molecular Biology and Evolution 14, 685–695 (1997). DOI: 10.1093/oxfordjournals.molbev.a025808.

13. Felsenstein, J. Evolutionary trees from DNA sequences: A maximum likelihood approach. Journal of Molecular Evolution 17, 368–376 (1981). DOI: 10.1007/BF01734359.

14. Parra, R. G. et al. Reconstructing complex lineage trees from scRNA-seq data using MER-LoT. Nucleic Acids Research 47, 8961–8974 (2019). DOI: 10.1093/nar/gkz706.

15. Garrido, Q. et al. Visualizing hierarchies in scRNA-seq data using a density tree-biased autoencoder. Bioinformatics 38, i316–i324 (2022). DOI: 10.1093/bioinformatics/btac249.

16. Yao, Z. et al. A high-resolution transcriptomic and spatial atlas of cell types in the whole mouse brain. Nature 624, 317–332 (2023). DOI: 10.1038/s41586-023-06812-z.

17. Moravec, J. C., Lanfear, R., Spector, D. L., Diermeier, S. D. & Gavryushkin, A. Testing for Phylogenetic Signal in Single-Cell RNA-Seq Data. Journal of Computational Biology 30, 518–537 (2023). DOI: 10.1089/cmb.2022.0357.

18. Mah, J. L. & Dunn, C. W. Cell type evolution reconstruction across species through cell phylogenies of single-cell RNA sequencing data. Nature Ecology & Evolution 8, 325–338 (2024). DOI: 10.1038/s41559-023-02281-9.

19. Domcke, S. & Shendure, J. A reference cell tree will serve science better than a reference cell atlas. Cell 186, 1103–1114 (2023). DOI: 10.1016/j.cell.2023.02.016.

20. Church, S. H., Mah, J. L. & Dunn, C. W. Integrating phylogenies into single-cell RNA sequencing analysis allows comparisons across species, genes, and cells. PLOS Biology 22, e3002633 (2024). DOI: 10.1371/journal.pbio.3002633.

21. Hao, Y. et al. Dictionary learning for integrative, multimodal and scalable single-cell analysis. Nature Biotechnology 42, 293–304 (2024). DOI: 10.1038/s41587-023-01767-y.

22. Becht, E. et al. Dimensionality reduction for visualizing single-cell data using UMAP. Nature Biotechnology 37, 38–44 (2019). DOI: 10.1038/nbt.4314.

23. Healy, J. & McInnes, L. Uniform manifold approximation and projection. Nature Reviews Methods Primers 4, 82 (2024). DOI: 10.1038/s43586-024-00363-x.

24. Minh, B. Q. et al. IQ-TREE 2: New Models and Efficient Methods for Phylogenetic Inference in the Genomic Era. Molecular Biology and Evolution 37, 1530–1534 (2020). DOI: 10.1093/molbev/msaa015.

25. Schwabe, D., Formichetti, S., Junker, J. P., Falcke, M. & Rajewsky, N. The transcriptome dynamics of single cells during the cell cycle. Molecular Systems Biology 16, e9946 (2020). DOI: 10.15252/msb.20209946.

26. Pavel, A., Grønberg, M. G. & Clemmensen, L. H. The impact of dropouts in scRNAseq dense neighborhood analysis. Computational and Structural Biotechnology Journal 27, 1278–1285 (2025). DOI: 10.1016/j.csbj.2025.03.033.

27. Linderman, G. C. et al. Zero-preserving imputation of single-cell RNA-seq data. Nature Communications 13, 192 (2022). DOI: 10.1038/s41467-021-27729-z.

28. Chen, J., Ng, Y. K., Lin, L., Zhang, X. & Li, S. On triangle inequalities of correlation-based distances for gene expression profiles. BMC Bioinformatics 24, 1–16 (2023). DOI: 10.1186/s12859-023-05161-y.

29. Huang, W. et al. TreeScaper: Visualizing and Extracting Phylogenetic Signal from Sets of Trees. Molecular Biology and Evolution 33, 3314–3316 (2016). DOI: 10.1093/molbev/msw196.

30. Jombart, T., Kendall, M., Almagro-Garcia, J. & Colijn, C. treespace: Statistical exploration of landscapes of phylogenetic trees. Molecular Ecology Resources 17, 1385–1392 (2017). DOI: 10.1111/1755-0998.12676.

31. Smith, M. R. Robust Analysis of Phylogenetic Tree Space. Systematic Biology 71, 1255–1270 (2022). DOI: 10.1093/sysbio/syab100.

32. Rubbi, A., Goldman, N. & Marleen, I. Pear-EBI (2024). URL https://github.com/AndreaRubbi/Pear-EBI.

33. Cannoodt, R., Saelens, W., Deconinck, L. & Saeys, Y. Spearheading future omics analyses using dyngen, a multi-modal simulator of single cells. Nature Communications 12, 3942 (2021). DOI: 10.1038/s41467-021-24152-2.

34. Gillespie, D. T. Exact stochastic simulation of coupled chemical reactions. The Journal of Physical Chemistry 81, 2340–2361 (1977). DOI: 10.1021/j100540a008.

35. Smith, M. R. Using Information Theory to Detect Rogue Taxa and Improve Consensus Trees. Systematic Biology 71, 1088–1094 (2022). DOI: 10.1093/sysbio/syab099.

36. Crowell, H. L., Morillo Leonardo, S. X., Soneson, C. & Robinson, M. D. The shaky foundations of simulating single-cell RNA sequencing data. Genome Biology 24, 1–19 (2023). DOI: 10.1186/s13059-023-02904-1.

37. Bastidas-Ponce, A. et al. Comprehensive single cell mRNA profiling reveals a detailed roadmap for pancreatic endocrinogenesis. Development 146, dev173849 (2019). DOI: 10.1242/dev.173849.

38. Fu, Q. et al. A comparison of scRNA-seq annotation methods based on experimentally labeled immune cell subtype dataset. Briefings in Bioinformatics 25, bbae392 (2024). DOI: 10.1093/bib/bbae392.

39. Zhang, Y. et al. Single-cell analyses of renal cell cancers reveal insights into tumor microenvironment, cell of origin, and therapy response. Proceedings of the National Academy of Sciences of the United States of America 118, e2103240118 (2021). DOI: 10.1073/p-nas.2103240118.

40. Traag, V. A., Waltman, L. & van Eck, N. J. From Louvain to Leiden: guaranteeing well-connected communities. Scientific Reports 9, 5233 (2019). DOI: 10.1038/s41598-019-41695-z.

41. Heumos, L. et al. Best practices for single-cell analysis across modalities. Nature Reviews Genetics 24, 550–572 (2023). DOI: 10.1038/s41576-023-00586-w.

42. Bryant, D. & Moulton, V. Neighbor-Net: An Agglomerative Method for the Construction of Phylogenetic Networks. Molecular Biology and Evolution 21, 255–265 (2004). DOI: 10.1093/molbev/msh018.

43. Bryant, D. & Huson, D. H. NeighborNet: improved algorithms and implementation. Frontiers in Bioinformatics 3 (2023). DOI: 10.3389/fbinf.2023.1178600.

44. Huson, D. H. & Bryant, D. The SplitsTree App: interactive analysis and visualization using phylogenetic trees and networks. Nature Methods 21, 1773–1774 (2024). DOI: 10.1038/s41592-024-02406-3.

45. Hejase, H. A. & Liu, K. J. A scalability study of phylogenetic network inference methods using empirical datasets and simulations involving a single reticulation. BMC Bioinformatics 17, 1–12 (2016). DOI: 10.1186/s12859-016-1277-1.

46. Philippe, H. & Lopez, P. On the conservation of protein sequences in evolution. Trends in Biochemical Sciences 26, 414–416 (2001). DOI: 10.1016/S0968-0004(01)01877-1.

47. Sanderson, M. J., McMahon, M. M., Stamatakis, A., Zwickl, D. J. & Steel, M. Impacts of Terraces on Phylogenetic Inference. Systematic Biology 64, 709–726 (2015). DOI: 10.1093/sysbio/syv024.

48. Huelsenbeck, J. P. & Ronquist, F. MRBAYES: Bayesian inference of phylogenetic trees. Bioinformatics 17, 754–755 (2001). DOI: 10.1093/bioinformatics/17.8.754.

49. Drummond, A. J. & Rambaut, A. BEAST: Bayesian evolutionary analysis by sampling trees. BMC Evolutionary Biology 7, 214 (2007). DOI: 10.1186/1471-2148-7-214.

50. Bouckaert, R. et al. BEAST 2: A Software Platform for Bayesian Evolutionary Analysis. PLOS Computational Biology 10, e1003537 (2014). DOI: 10.1371/journal.pcbi.1003537.

51. De Maio, N. et al. Maximum likelihood pandemic-scale phylogenetics. Nature Genetics 55, 746–752 (2023). DOI: 10.1038/s41588-023-01368-0.

52. Charif, D. & Lobry, J. R. in SeqinR 1.0-2: A Contributed Package to the R Project for Statistical Computing Devoted to Biological Sequences Retrieval and Analysis (eds Bastolla, U., Porto, M., Roman, H. E. & Vendruscolo, M.) Structural Approaches to Sequence Evolution: Molecules, Networks, Populations 207–232 (Springer, Berlin, Heidelberg, 2007).

53. Kalyaanamoorthy, S., Minh, B. Q., Wong, T. K. F., von Haeseler, A. & Jermiin, L. S. ModelFinder: fast model selection for accurate phylogenetic estimates. Nature Methods 14, 587–589 (2017). DOI: 10.1038/nmeth.4285.

54. Yu, G. Using ggtree to Visualize Data on Tree-Like Structures. Current Protocols in Bioinformatics 69, e96 (2020). DOI: 10.1002/cpbi.96.

55. Csardi, G. & Nepusz, T. The igraph software. Complex syst 1695, 1–9 (2006).

56. Bergen, V., Lange, M., Peidli, S., Wolf, F. A. & Theis, F. J. Generalizing RNA velocity to transient cell states through dynamical modeling. Nature Biotechnology 38, 1408–1414 (2020). DOI: 10.1038/s41587-020-0591-3.

57. Fu, Q. A Comparison of scRNA-seq Annotation Methods Based on Experimentally Labeled Immune Cell Subtype Dataset (2024). URL https://zenodo.org/records/10947879.

